# Cytoplasmic forces functionally reorganize nuclear condensates in oocytes

**DOI:** 10.1101/2021.03.15.434387

**Authors:** Adel Al Jord, Gaëlle Letort, Soline Chanet, Feng-Ching Tsai, Christophe Antoniewski, Adrien Eichmuller, Christelle Da Silva, Jean-René Huynh, Nir S. Gov, Raphaël Voituriez, Marie-Émilie Terret, Marie-Hélène Verlhac

## Abstract

Cells remodel their cytoplasm with force-generating cytoskeletal motors^1^. Their activity generates random forces that stir the cytoplasm, agitating and displacing membrane-bound organelles like the nucleus in somatic^2–4^ and germ^5–7^ cells. These forces are transmitted inside the nucleus^4,7^, yet their consequences on liquid-like biomolecular condensates^8–10^ residing in the nucleus remain unexplored. Here, we probe experimentally and computationally diverse nuclear condensates, that include nuclear speckles, Cajal bodies, and nucleoli, during cytoplasmic remodeling of female germ cells named oocytes. We discover that growing mammalian oocytes deploy cytoplasmic forces to timely impose multiscale reorganization of nuclear condensates for the success of meiotic divisions. These cytoplasmic forces accelerate nuclear condensate collision-coalescence and molecular kinetics within condensates. Inversely, disrupting the forces decelerates nuclear condensate reorganization on both scales, compromising condensate-associated mRNA processing and consequently hindering oocyte divisions that drive female fertility. We establish that cytoplasmic forces can reorganize nuclear condensates in an evolutionary conserved fashion in insects. Our work implies that cells evolved a mechanism, based on cytoplasmic force tuning, to functionally regulate a broad range of nuclear condensates across scales. This finding opens new perspectives when studying condensate-associated pathologies like cancer, neurodegeneration and viral infections^11–13^.

## Main text

Fertility depends on essential maternal transcripts and proteins accumulated during growth of developing female germ cells known as oocytes^14^. In mammals, growing oocytes remodel their cytoplasm to position the nucleus at the cell center despite undergoing an asymmetric division in size, which relies on chromosome off-centering, soon after^15–17^. Central nucleus position predicts successful oocyte development in mice and humans^15,16^, and thus, their embryogenic potential. However, the spatiotemporal evolution and the overall functional role of this cytoplasmic remodeling remain unclear.

Cytoplasmic remodeling is driven by cytoskeletal motor proteins^1^. Motor activity generates random fluctuating forces that stir the cytoplasm, enhancing nonspecific transport of organelles and their agitation^2–7^. In growing mouse oocytes, cytoplasmic random stirring is predominantly driven by F-actin and myosin-motor activity^5^ with physical consequences observable on two spatiotemporal scales. On a large scale, prolonged cytoplasmic stirring displaces, nonspecifically, the 30µm nucleus from cell periphery to its center over a 25µm distance^5,18^. On a smaller scale, cytoplasmic stirring agitates the nucleus with stochastic kicks, resulting in nuclear membrane fluctuations and cytoplasmic force propagation inside the nucleus^6,7^. These forces enhance the mobility of the nucleolus^7^, one of many nuclear liquid-like compartments of RNA processing known as biomolecular condensates^8–10^. We observed that, as growing mouse oocytes remodeled their cytoplasm, the number of nucleoli decreased in favor of a size increase (Fig.1a-c), as noted long ago in human oocytes^19^. Moreover, fully grown oocytes mutant for the F-actin nucleator Formin2 and consequent capacity to stir their cytoplasm presented a peripheral nucleus enriched in multiple immobile nucleoli of smaller sizes (Fig.1a-c, Extended Data Fig.1a and (^7^)). Since nucleoli as other liquid condensates can coalesce, this prompted us to investigate whether cytoplasmic forces during oocyte growth orchestrated the reorganization of nuclear liquid condensates.

**Fig. 1.**
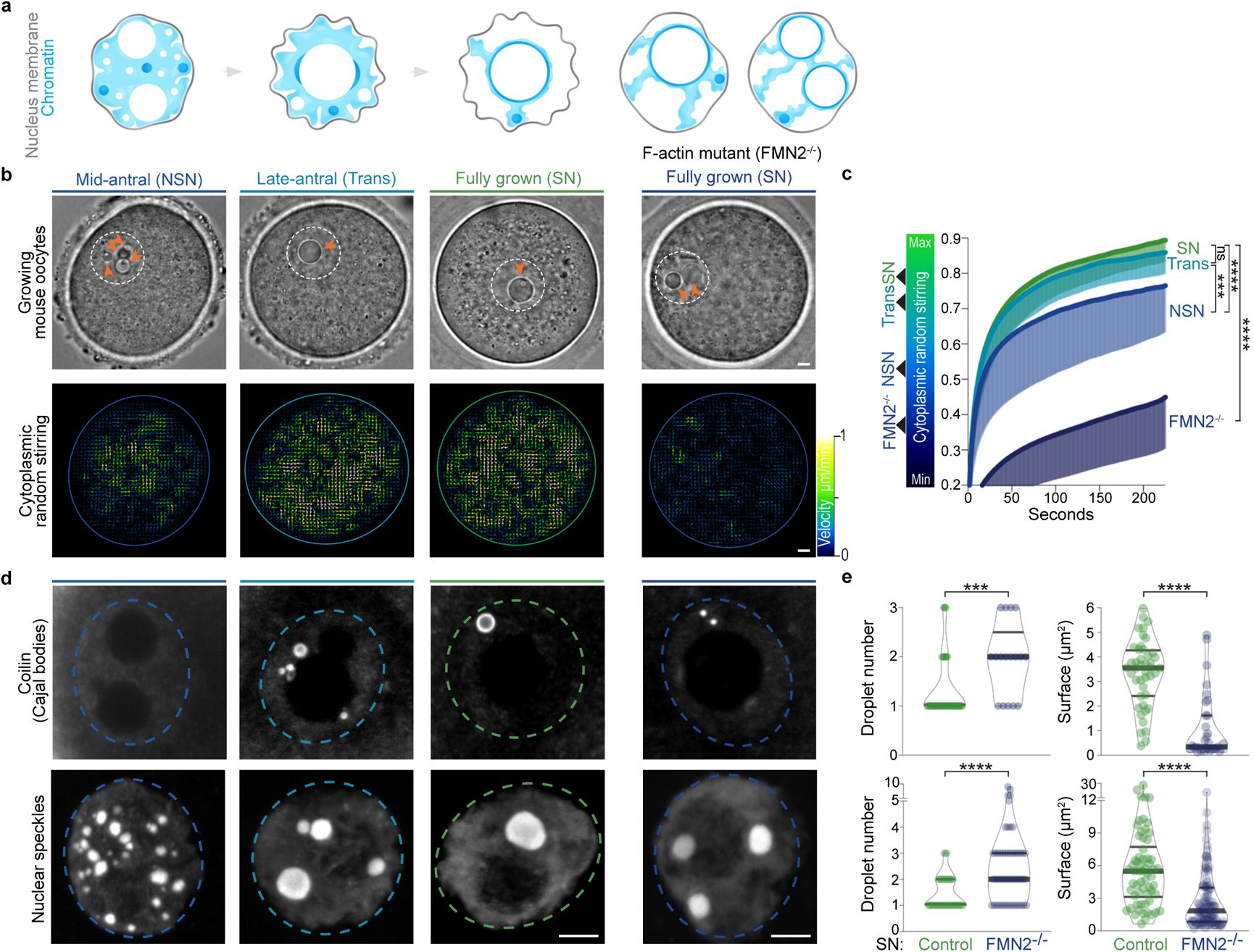
Nuclear biomolecular condensate patterns during mouse oocyte growth correlate with cytoplasmic remodeling. (**a**) Illustration of chromatin configuration evolution (NSN-Trans-SN) coinciding with oocyte consecutive growth subcategories (mid-antral follicle, late-antral follicle, and fully grown); (left) large spherical nucleoli decrease in number in favor of a size increase as of the Trans-stage; (right) chromatin configuration scenarios in fully grown cytoplasmic F-actin mutants (FMN2^−/−^) with one large or two smaller nucleoli. The nuclear membrane (gray) is depicted according to its fluctuation intensity at distinct stages of growth (see (^7^) and ED Fig.1j). (**b**) Top: Bright-field images of (left panels) Control oocytes with growth progression and (right panel) a fully grown mutant; the nucleus is outlined in dashed white and multiple small nucleoli (NSN and FMN2^−/−^ SN) or a single large nucleolus (Trans and SN) are indicated with orange arrowheads. Bottom: Growth-associated cytoplasmic stirring vector maps generated by STICS analyses of bright-field 240 seconds-stream videos; the oocyte cortex is outlined with colors reflecting the intensity of cytoplasmic stirring; maps are color coded according to velocity magnitude (lowest in dark blue, strongest in white). (**c**) Cytoplasmic random stirring intensity in time measured in Control NSN to SN and FMN2^−/−^ SN oocytes by image correlation analyses of cytoplasmic pixel evolution; color gradient (left) used in this study to represent the cytoplasmic stirring intensity; cell number, Control NSN=22, Trans=19, SN=28 cells, FMN2^−/−^ SN=13; error bars represent mean-s.d.; *P* values derived from two-tailed Mann-Whitney *U*-Tests on plateau distributions, ns, not significant, *P*=0.074, \*\*\**P*=0.0001, \*\*\*\**P*<0.0001. (**d**) Representative immunoreactivities of Coilin and nuclear speckles in growing Control oocytes and SN FMN2^−/−^ oocytes; Coilin images are 0.5µm z-planes and nuclear speckle images are 20µm z-projections; nucleus regions outlined with dashed circles. (**e**) Quantifications of Coilin (top) and nuclear speckle (bottom) droplet number and surface in nuclei of fully grown SN Control and mutant oocytes; Coilin, counted in 43 Control and 17 FMN2^−/−^ cells, measured 44 Control and 33 FMN2^−/−^ droplets; speckles, counted in 41 Control and 52 FMN2^−/−^ cells, measured 62 Control and 112 FMN2^−/−^ droplets; violin plots with median±quartiles; *P* values derived from two-tailed Mann-Whitney *U*-Tests, \*\*\**P*=0.0004, \*\*\*\**P*<0.0001. Color codes based on cytoplasmic stirring intensities; scale bars, 5µm.

To proceed, we first revised an oocyte late-growth classification method, solely based on progressive chromatin condensation^20^, by linking it to a measure of cytoplasmic stirring and consequent nucleus position (Fig.1a-c, Extended Data Fig.1b-k; Video 1). Growing oocytes extracted from antral follicles presented the three expected chromatin states in a nucleus of constant volume. Smaller mid-antral “NSN” oocytes (<u>N</u>on-<u>S</u>urrounded <u>N</u>ucleolus) had relatively low cytoplasmic F-actin and stirring with an off-centered poorly agitated nucleus, mirrored by moderate nucleus membrane fluctuations. Denser cytoplasmic F-actin and stronger cytoplasmic stirring were quantified in subsequent bigger late-antral “Trans” oocytes (<u>Trans</u>itioning) that were in the process of nucleus centering. Comparably dense F-actin and equally intense stirring were measured in the cytoplasm of fully grown “SN” oocytes (<u>S</u>urrounded <u>N</u>ucleolus), with high cytoplasmic forces propagated onto a centrally positioned nucleus with a strongly fluctuating nuclear membrane. Thus, oocytes maximally intensify cytoplasmic stirring as of the Trans growth-stage. This updated three-stage classification method, connecting known oocyte chromatin features with cytoplasmic stirring intensity, enabled us to inspect whether this cytoplasmic activity affects nuclear condensates in growing oocytes.

To explore a global relation between cytoplasmic stirring and nuclear liquid condensates, we screened for diverse nuclear biomolecular condensates^9^ in growing oocytes using known immunomarkers^21^ of histone locus bodies, paraspeckles, gems, nucleoli, TDP-43 bodies, Cajal bodies and nuclear speckles. Careful 3D-examination of these condensates revealed their nuclear pattern evolution from the NSN to SN stages (Extended Data Fig.2a-b). Further analyses demarcated condensates into two subpopulations with different properties. A first subpopulation of small shape-constrained condensates with stable chromatin association, insensitivity to the droplet dissolver 1-6-hexanediol^22^, and incomplete fluorescence recovery after photobleaching (Extended Data Figs.2c-d, 3a-h), collectively indicative of a non-liquid nature. A second subpopulation of large spherical and mobile droplet condensates that lacked apparent chromatin association, dissolved in response to 1-6-hexanediol, fully recovered fluorescence post-bleaching, and rapidly coalesced (Extended Data Figs.2c-d,3a-i; Videos 2-3), collectively indicative of liquid-like nature. We therefore focused on this second subpopulation of condensates to address our study’s key question. We counted and measured various nuclear droplets in growing Control oocytes and in growing Formin2-mutant oocytes (FMN2^−/−^), that present a significant drop in cytoplasmic stirring (Fig.1d-e; Extended Data Fig.4a-c). In Controls, Coilin (Cajal body), nuclear speckle, nucleolar, and TDP-43 droplets decreased in number and increased in size as growth ended. FMN2^−/−^ oocytes, however, had more numerous yet smaller Coilin, nuclear speckle, nucleolar, and TDP-43 droplets as they reached the fully-grown stage, despite Control-like total nuclear amounts of corresponding markers at tested growth stages (Extended Data Fig.4d-e). Other parameters known to modulate droplet size like cell and nucleus volume or neo-transcription were unaffected in mutants (Extended Data Fig.4f-j). Thus, the intensification of cytoplasmic stirring in growing oocytes is linked to the spatiotemporal evolution of several nuclear liquid condensates, potentially through increased droplet coalescence dynamics.

To confirm that cytoplasmic stirring enhanced nuclear droplet dynamics, we live-imaged at high temporal resolution growing oocytes microinjected with RNA encoding the nuclear speckle marker SRSF2-GFP. We chose speckles due to droplet abundance at all stages of growth and verified that SRSF2-GFP expression profiles were comparable to endogenous speckles. In all three stages of growth, we observed local droplet displacements and fusions instigated by the fluctuating nuclear membrane (Fig.2a, Extended Data Fig.5a; Video 3), an immediate consequence of cytoplasmic stirring forces agitating the nucleus^7^. To estimate the magnitude of these local cytoplasmic forces agitating the nuclear membrane, we optically trapped cytoplasmic vesicles and pushed them against the membrane, artificially inducing comparable nuclear invaginations and droplet displacements with a force of the order of 10pN (Fig.2b-d), in agreement with forces generated by multiple cytoskeletal motors in the oocyte cytoplasm^6^. Moreover, droplet diffusion was inversely proportional to its radius and correlated positively with the intensification of cytoplasmic forces during growth (Extended Data Fig.5b-c). This was consistent with our hypothesis that cytoplasmic forces induce active diffusion of nuclear droplets (see Biophysical model in Methods). We therefore tuned the transmission of cytoplasmic forces to the nucleoplasm, using established pharmacological and genetic tools^5,7^, before monitoring droplet diffusion at distinct growth stages (Fig.2e-f, Extended Data Fig.5d-e; Video 4). Amplifying nucleus agitation by disrupting force-dampening microtubules with Nocodazole^7^ increased the effective diffusion of droplets at all stages when compared to Controls. Inversely, obstructing forces from the nucleus by Taxol-mediated stabilization of microtubules slowed down droplets. Greater deceleration occurred in cells with disrupted cytoplasmic actomyosin-based forces due to Cytochalasin-D incubation or FMN2 knockout. Depleting residual microtubule-based forces in FMN2^−/−^ oocytes with Nocodazole further diminished nuclear droplet diffusion. Droplet diffusion generally remained sub-diffusive (Extended Data Fig.5f; *α*<1), consistent with chromatin-mediated constraints on droplet diffusion^23^. Thus, cytoplasmic forces, by agitating the nucleus, enhance diffusive dynamics of nuclear droplets in constrained environments. Moreover, tuning nucleus agitation led to droplet size changes matching agitation intensity without affecting droplet shape or total SRSF2-GFP levels (Extended Data Fig.5g-i). This suggested that cytoplasmic stirring, by enhancing droplet diffusivity, favored nuclear droplet encounters and coalescence.

**Fig. 2.**
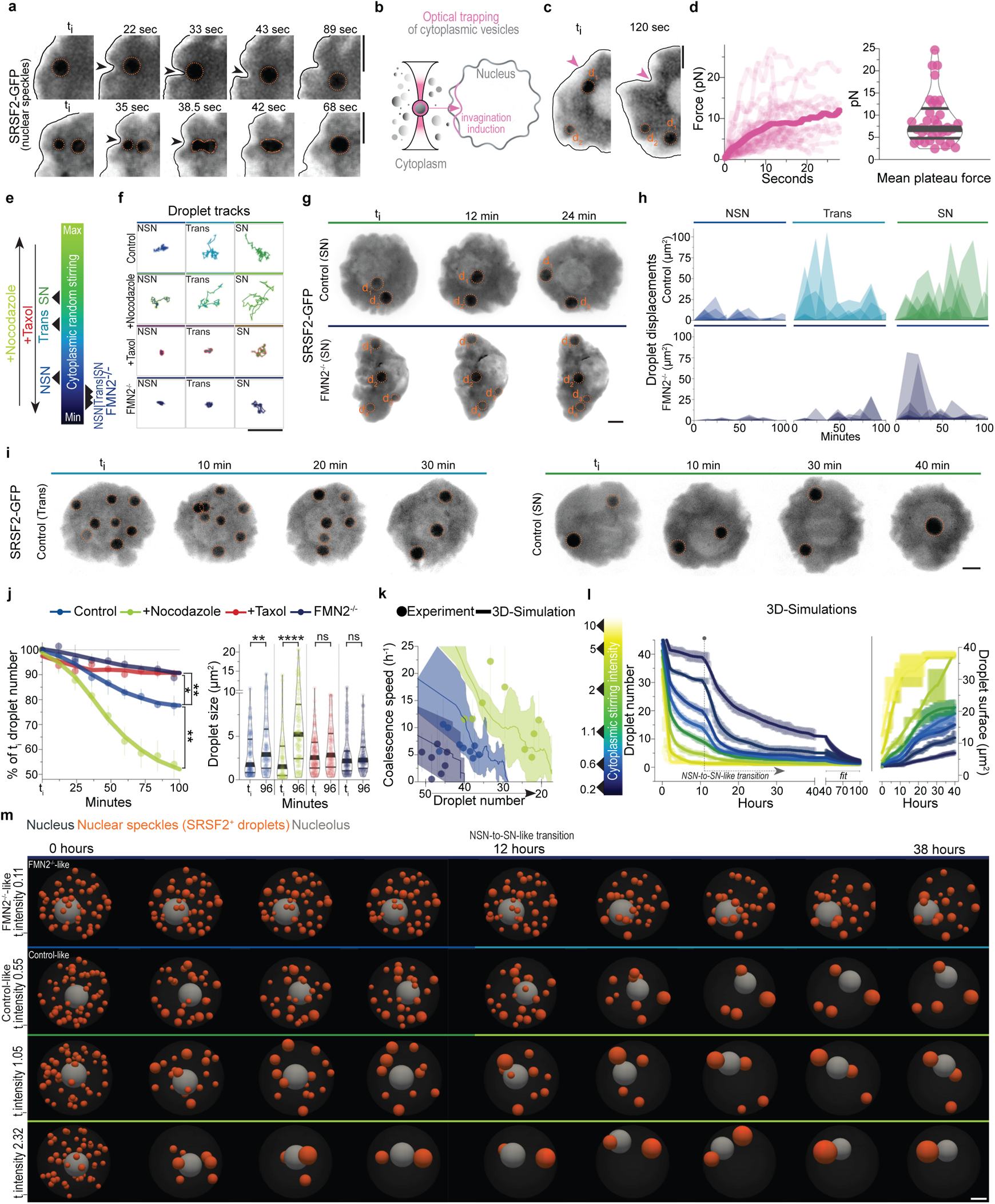
Cytoplasmic forces drive mesoscale condensate collision-coalescence in the nucleus. (**a**) Time-lapse 2D-images showing nuclear membrane invaginations physically kicking and locally displacing nucleoplasmic SRSF2-GFP droplets (seconds timescale) in Trans (top) or NSN (bottom) cells, directly causing droplet collision-coalescence; black arrowheads indicate nuclear membrane invaginations and the membrane is outlined in black. (**b**) Illustration of optical trapping experiment; cytoplasmic vesicles close to the nucleus were optically trapped and pushed against the nuclear membrane to exert forces upon it and induce nuclear membrane invaginations. (**c**) Time-lapse images showing a nuclear membrane invagination induced by the optical trapping approach and displacing a nucleoplasmic SRSF2-GFP droplet in a Trans cell; pink arrowhead indicates point of contact between trapped cytoplasmic vesicle and nuclear membrane. (**d**) Individual force tracks in time (left) and mean plateau forces (right) obtained with the optical trapping approach; 37 measurements from 20 oocytes. (**e**) Color gradient used in this study to represent the intensity of cytoplasmic forces in growing Control and mutant oocytes or Controls treated with microtubule inhibitors; Nocodazole disrupts perinuclear force-dampening by microtubules thus intensifying force transmission, whereas Taxol stabilizes microtubules, obstructing cytoplasmic forces from the nucleus. (**f**) Representative SRSF2-GFP droplet 2D-space exploration on a seconds-timescale (50 second-tracks) at the three growth stages in Control oocytes, Controls incubated with Nocodazole or Taxol, and FMN2^−/−^ oocytes. (**g**) Time-lapse 50µm z-projections showing minute-timescale 3D-displacements of tracked SRSF2-GFP droplets in the nucleoplasm of fully-grown SN Control and FMN2^−/−^ oocytes. (**h**) 3D-tracks of SRSF2-GFP droplets revealing fluctuating displacements (squared) in Control and FMN2^−/−^ oocytes at three stages of growth; droplet number in Control, NSN=8, Trans=5, SN=6; in FMN2^−/−^, NSN=12, Trans=11, SN=26. (**i**) Control SRSF2-GFP droplet minute-scale 3D-dynamics (50µm z-projections) in Trans and SN cells highlighting random nucleoplasmic displacements and collision-coalescence dynamics. (**j**) Nuclear SRSF2-GFP droplet collision-coalescence speed at the minute to hour-timescales. (Left) nuclear SRSF2-GFP droplet number evolution (in % of initial droplet number) with Control, amplified (+Nocodazole), or disrupted cytoplasmic forces (+Taxol or FMN2^−/−^) in NSN oocytes; oocyte number, 4 oocytes for each condition; error bars represent mean±s.e.m.; *P* values derived from two-tailed Wilcoxon matched-pairs signed rank tests, \**P*=0.0195, \*\**P*<0.0078. (Right) nuclear droplet surface evolution between the initial timepoint (t_i_) and t=96 min; droplet number, Control t_i_=113,t_96_=96, Nocodazole t_i_=93,t_96_=58, Taxol t_i_=114,t_96_=104, FMN2^−/−^ t_i_=141,t_96_=124; violin plots with median±quartiles; *P* values derived from two-tailed Mann-Whitney *U*-Tests, ns, not significant, *P*>0.177, \*\**P*=0.0071, \*\*\*\**P*<0.0001. (**k**) Nuclear droplet coalescence speed per hour relative to droplet number decrease in experimental NSN conditions (Control in blue, Nocodazole in green and FMN2^−/−^ in dark blue) and computational models with simulated NSN Control-like, Nocodazole-like, and FMN2^−/−^-like cytoplasmic activity; error represents mean±95% confidence interval; number of experimental measurements, Control NSN=9, Nocodazole=9, FMN2^−/−^=7; 5 simulations for each condition. (**l**) 3D-simulations (n=41) showing nucleoplasmic SRSF2 droplet coalescence speed on an hour-to-day timescale relative to cytoplasmic forces (intensities colored according to the gradient on the left); simulations start from an NSN-like nucleus state with 45 SRSF2 droplets; chromatin condensation and cytoplasmic activity intensification, simulating the transition into late growth, occurs at 12 hours. (**m**) Time frames of 3D-simulations in (l) with a gradient of starting-point cytoplasmic intensities that include the FMN2^−/−^-like and Control-like scenarios; nuclear speckles (SRSF2^+^ droplets) are in orange and a single grey nucleolus is depicted in a spherical nucleus-like container. Droplets in **a**, **c, g**, and **i** are outlined in dashed orange; color codes based on cytoplasmic stirring intensities; scale bars, 5µm.

To verify the cytoplasmic drive of nuclear droplet coalescence, we documented longer-term consequences of cytoplasmic stirring forces on nuclear droplet dynamics. 3D-tracking revealed stochastic displacements of droplets that bounced randomly around the nucleoplasm more vigorously as of the Trans-stage (Fig.2g-h), concomitant with cytoplasmic stirring intensification. Droplets eventually encountered one another, collided and fused (Fig.2i, Extended Data Fig.6a; Video 5). In FMN2 mutants, droplets bounced less frequently, and this decreased further when both F-actin and microtubules were disrupted (Fig.2g-h, Extended Data Fig.6b-d). We therefore modulated cytoplasmic forces and measured nuclear droplet coalescence kinetics in NSN oocytes that harbor significantly more droplets than SN counterparts (Fig.2j, Extended Data Fig.6e-g; Video 6). Amplifying cytoplasm-based nucleus agitation accelerated droplet coalescence, manifested by a quicker decline in droplet number coupled to droplet size increase when compared to Controls. Inversely, taming nucleus agitation decelerated droplet coalescence, revealed by weak droplet number decline and mean droplet size changes within the 100-minute filming period. Chromatin remained in a decondensed state in analyzed cells (Extended Data Fig.6h-i), hence eliminating the chromatin condensation bias potentially impacting droplet coalescence. Thus, cytoplasm-based agitation of the nucleus accelerates nuclear droplet collision-coalescence kinetics.

To further dissect the cytoplasmic drive of nuclear droplet collision-coalescence, we built an agent-based model of SRSF2-like droplet diffusion in a nucleus-like spherical container agitated by cytoplasmic forces. Nuclear obstacles, based on experimental chromatin surface measurements, were added to mimic nuclear crowding by chromatin. 3D-numerical simulations were calibrated to recapitulate Control-NSN nuclear droplet diffusive dynamics on the seconds timescale (Extended Data Fig.7a). Unexpectedly, running these simulations on 100-minute timescales reproduced experimental Control and modulated cytoplasmic drives of nuclear droplet coalescence in NSN oocytes (Fig.2k, Extended Data Fig.7b). Thus, nuclear droplet diffusion and collision-coalescence propelled by cytoplasmic forces are sufficient to explain the observed droplet coarsening in the crowded nucleus.

To then estimate the minimal time necessary to reorganize nuclear compartments if cells were to transition from NSN to SN states of oocyte growth, a process occurring inside the follicle that cannot be followed in live, and to evaluate chromatin’s interference with the process, we ran our computational model on longer timescales (Fig.2l-m, Extended Data Fig.7c-e; Video 7). To reproduce key physiological cytoplasmic and nucleoplasmic events that initiate at the end of oocyte growth, we incorporated into our simulations force intensification and chromatin condensation (Simulation series 2 with transition at 12 hours) that occurs after the NSN-stage. To reach the endogenous compartmentalization state of 4 nuclear speckle droplets observed at the final growth stage (mean speckle droplet number in Trans cells with a central nucleus), Control oocytes would require 15±8 hours from the NSN|Trans transition. This prediction is consistent with *in vivo* mid-to-end of antral oocyte growth, roughly estimated to occur within less than 2 days physiologically. The prediction is also consistent with prior experimental *in vitro* and computational studies since it falls within the time-range of 5 to 17 hours necessary for cytoplasmic activity to displace the peripheral nucleus to the cell center^5,18^. By contrast, cells with diminished forces insufficient to transport the nucleus, such as FMN2^−/−^ oocytes^5,18^, would require at least 59±8 hours to reach a nuclear compartmentalization state comparable to Controls. These simulations also highlighted chromatin condensation-based acceleration of droplet fusion (compare before|after t=12h in Fig.2l), which would rationalize experimental observations of faster droplet diffusion in mutants that initiate chromatin condensation without intensifying cytoplasmic forces (Extended Data Fig.5e). We therefore generated two supplementary models to weigh chromatin’s interference with nuclear droplet kinetics (Series 1 & 3). We modelled an NSN nucleus state with decondensed chromatin and a SN nucleus state with condensed chromatin before launching simulations with force intensities maintained constant in time. Droplet collision-coalescence was faster in the condensed chromatin set-up (Series 3), suggesting a kinetic barrier effect for chromatin coherent with its solid-like and diffusion-constraining properties^23,24^. Probing further, we found that with weak forces (FMN2^−/−^-like scenario), chromatin significantly hindered droplet fusion, seen by the substantial drop in time required to coalesce all droplets post-chromatin condensation (Extended Data Fig.7f; compare FMN2^−/−^ plain and dashed fits). In all Control-like scenarios, however, the forces reduced chromatin’s hindrance of droplet fusion (Extended Data Fig.7f; compare NSN, Trans, and SN plain and dashed fits). Thus, our simulations anticipate cytoplasmic force intensification to be a predominant driver of nuclear droplet coalescence in physiological contexts. Experiments and computations indicate that cytoplasmic forces in growing oocytes spatially reorganize nuclear liquid compartments by boosting large-scale droplet collision-coalescence.

To determine local sub-droplet-scale consequences of cyto-to-nucleoplasmic force transfer, we first measured rapid surface fluctuations of large SRSF2-GFP droplets in Control and F-actin mutant SN oocytes±Nocodazole. Surface fluctuations of comparably-sized droplets declined when F-actin-based cytoplasmic forces were disrupted, and decreased further after simultaneous disruption of both F-actin and microtubules (Fig.3a, Extended Data Fig.8a). The Control:mutant droplet fluctuation ratio (≅2) was smaller than the Control:mutant nuclear membrane fluctuation ratio (≅6 in (^7^)), in agreement with expected energy dissipation theorized by our biophysical model linking cytoplasmic stirring with the nucleoplasm (see Methods). This observable energy transfer of cytoplasmic forces onto nuclear droplets prompted us to evaluate the potential enhancement of droplets’ internal molecular kinetics using FRAP (Fig.3b-c, Extended Data Fig.8b-e). Droplet molecules exchanged significantly slower in FMN2^−/−^ cells and cells treated with Cytochalasin-D than in Controls. Molecular dynamics in droplets were reversible after restoration of cytoplasmic stirring by drug washout. To quantify the contribution of cytoplasmic active forces in driving nuclear dynamics across scales, we estimated a Péclet number P_n_ at the scale of droplet diffusion in the nucleoplasm down to molecular diffusion in the nucleoplasm and within droplets. This number can be defined here as the ratio of the expected diffusion coefficient resulting from active forces only, inferred from kinetic measurements of active fluctuations of the nuclear membrane and droplet surface, to the observed diffusion coefficient that combines the response to both active stirring and thermal diffusion. Our calculations indicate that the active component is dominant in driving the documented dynamics - from diffusion of droplets in the nucleoplasm (P_n_~5) down to that of molecules in the nucleoplasm (P_n_~2.5) and within droplets (P_n_~1.5) - as the energy of cytoplasmic active forces cascades across scales (see Biophysical model in Methods). Thus, cytoplasm-based agitation of the nucleus enhances nuclear sub-droplet-scale kinetics by amplifying droplet surface fluctuations and interior molecular mobility.

**Fig. 3.**
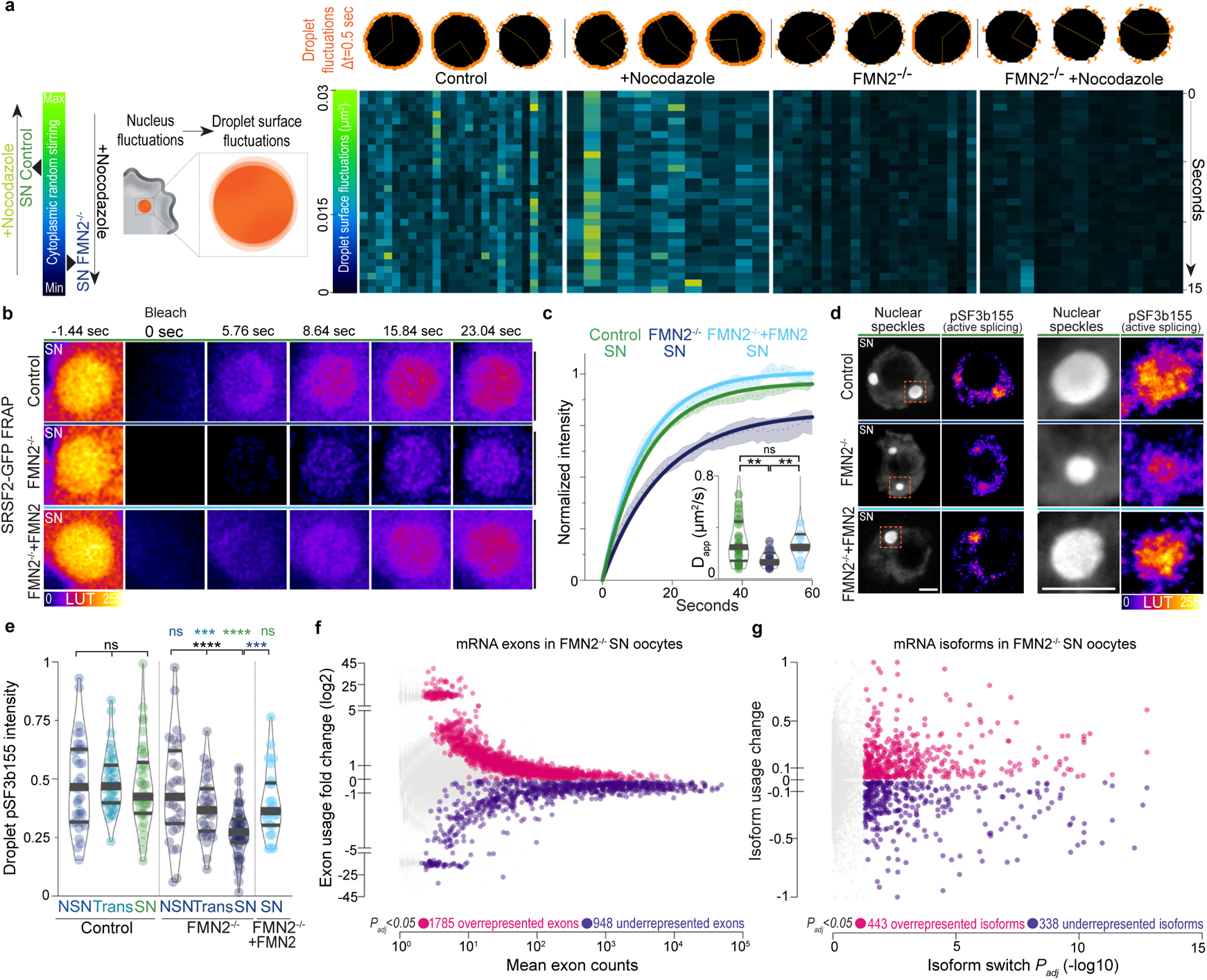
Cytoplasmic forces accelerate molecular-scale kinetics in nuclear condensates and regulate mRNA processing. (**a**) Scheme of droplet surface fluctuations as a function of cytoplasmic stirring (left); representative nuclear SRSF2-GFP droplet surface fluctuations (above right, droplet surface fluctuations in orange and droplet interiors are masked in black) and surface fluctuation intensity heatmaps of comparably sized droplets (below right; each column corresponds to a droplet and rows correspond to time) in SN oocytes with Control, amplified, or disrupted cytoplasmic forces; range of droplet radii, 2 to 2.7µm; droplet number, Control=25; Nocodazole=12; FMN2^−/−^=21; FMN2^−/−^+Nocodazole=15. (**b**) Representative FRAP sequences of SRSF2-GFP droplets in Control, FMN2^−/−^, and FMN2^−/−^+FMN2 SN oocytes. (**c**) Normalized fluorescence intensity recovery curves (mean±s.e.m.) with simple exponential fits of SRSF2-GFP droplets in Control, FMN2^−/−^, and FMN2^−/−^+FMN2 SN oocytes; insets, apparent diffusion coefficients (*D_app_*); droplet number, Control=28, FMN2^−/−^=13, FMN2^−/−^+FMN2=11; violin plots with median±quartiles; *P* values derived from two-tailed Mann-Whitney *U*-Tests, ns, not significant, *P=*0.7939, \*\**P*<0.0099. (**d**) Representative co-immunostainings of nuclear speckles and phosphorylated SF3b155 (pT313; active splicing marker) in SN Control, FMN2^−/−^, and FMN2^−/−^+FMN2 oocytes with higher magnifications of single droplets. (**e**) Quantifications of single droplet-specific pT313-SF3b155 intensities in Control and FMN2^−/−^ oocytes at the three growth stages, and in FMN2^−/−^+FMN2 SN oocytes; Droplet number, Control NSN=34, Trans=37, SN=38, FMN2^−/−^ NSN=31, Trans=33, SN=56, FMN2^−/−^+FMN2 SN=26; violin plots with median±quartiles; *P* values derived from two-tailed Mann-Whitney *U*-Tests and Kruskal-Wallis Tests, ns, not significant, *P*>0.0678, \*\*\**P*<0.0006, \*\*\*\**P*<0.0001. (**f-g**) Differential mRNA exon usage (log2) versus mean abundance (mean exon counts) or differential mRNA isoform usage versus isoform switch *P_adj_* (-log10) in SN FMN2^−/−^ oocytes relative to Control SN FMN2^+/−^ oocytes obtained with RNA-sequencing (from (^7^)) and DEXSeq or IsoformSwitchAnalyzeR packages; Colored dots are significantly over/underrepresented (fuchsia/purple) exons or isoforms in FMN2^−/−^ oocytes with a BH-adjusted (exons) or FDR-adjusted (isoforms) *P* value *P_ad_*_j_<0.05. Scale bars, 5µm.

Biomolecule mobility in condensates is expected to boost local biochemical reactions, which in nuclear speckles corresponds to, but is not limited to, mRNA processing by the spliceosome^25,26^. RNA-binding splicing regulators like the SR-protein SRSF2 activate and alter splicing by binding specific regions of pre-mRNA known as exon splicing enhancers^27^. We therefore hypothesized that cytoplasmic forces, by accelerating molecular dynamics in droplets, could enhance protein-RNA interaction kinetics that regulate splicing. To first test the splicing activation hypothesis, we probed catalytically active spliceosomes engaged in mRNA splicing using a specific phospho-marker (pThr313-SF3b155)^25^ during oocyte growth ±cytoplasmic stirring. Consistent with the general transcriptional decline during Control and mutant growth (Extended Data Fig.4j), total nucleoplasmic splicing activity assessed with pSF3b155 decreased while total unphosphorylated SF3b155 remained comparable (Extended Data Fig.8f-i). As opposed to the comparable transcriptional decline in both contexts, total nucleoplasmic splicing in mutants decreased more than in Controls as of the Trans stage (Extended Data Fig.8g). Since splicing was either droplet-associated or, when apparently nucleoplasmic, mainly chromatin-associated (Extended Data Fig.8j), we compared droplet-associated to chromatin-associated splicing ratios in Controls and mutants. Along with the transcriptional decline, splicing activity switched to primarily droplet-associated splicing in both SN Controls and mutants (Extended Data Fig.8k), while residual chromatin-associated splicing remained comparable (Extended Data Fig.8l). This implied that the primary difference in splicing was occurring in nuclear droplets, prompting us to compare droplet-associated splicing in growing Control and mutant oocytes. Active splicing remained unexpectedly constant in Control droplets while it dropped in droplets of mutants as of the Trans-stage (Fig.3d-e). Since cytoplasmic stirring in Controls intensifies as of the Trans-stage, this suggested that cytoplasmic force-intensification sustains splicing activity specifically in nuclear droplets.

To next test the alternative splicing hypothesis, we performed splicing-centered bioinformatic analyses of mRNA-sequencing data comparing Control with mutant SN oocytes^7^, the stage during which the observed splicing decline was maximal. We detected thousands of exon usage differences in mutant oocytes with 1785 overrepresented and 948 underrepresented exons (Fig.3f; Table 1), suggesting alternative splicing with enrichment of exon inclusion that is compatible with the idea of disrupted SR protein-RNA binding^28^. To probe the mRNA-processing imbalance further, we scrutinized differential transcript isoform usage, a direct consequence of alternative splicing, and uncovered hundreds of isoform switches in mutant oocytes (Fig.3g; Table 2). Multiple differential splicing events per transcript were identified for a sum of 3565 alternative splicing patterns of different subtypes in 1259 transcripts, with 2178 predicted consequences on transcript length or stability (Extended Data Fig.9a-c). We confirmed some splicing patterns with RT-qPCR (Extended Data Fig.9d-f). Supplementary enrichment and spatial correlation tests revealed that: altered exons and transcripts were significantly enriched in SRSF1 and SRSF2 binding sites; and altered splicing sites significantly overlapped with SRSF1 and SRSF2 sites up to 35 times more than other RNA-binding proteins, including the muscle-specific splicing factor MBNL3^29^ or the RNA-binding transcription factor YY1^30^ (Extended Data Fig.9g; Table 3). Consistent with our hypothesis, these data imply that cytoplasmic forces enhance SR protein-RNA binding kinetics in nuclear droplets, and perhaps the binding kinetics of other splicing regulators nearby. To finally confirm the mechanistic link between cytoplasmic forces and splicing, we expressed FMN2 in FMN2^−/−^ SN oocytes to restore cytoplasmic stirring. In these oocytes with rescued forces, multiscale kinetics of nuclear droplets were restored as well as their number and size within 5 hours (Fig.3b-c, Extended Data Fig.9h-n). Splicing activity was also fully restored in droplets without affecting residual chromatin-associated splicing (Fig.3d-e, Extended Data Fig.9o). Splicing patterns were reversed to a certain extent with a mean rescue efficiency of ~60% (Extended Data Fig.9d-f; compare rescue fold changes to mutant ones), which was expected due to accumulation of mis-spliced transcripts prior to the rescue. Thus, cytoplasmic forces regulate mRNA processing by enhancing droplet-associated splicing in growing oocytes with potentially broad consequences on maternal reservoir transcripts.

To seize the developmental impact of cytoplasmic force-based regulation of splicing, we analyzed the functional enrichment of altered transcripts in FMN2-mutants. Genes concerned with mRNA-processing alterations were enriched for diverse cell division-related functions (Table 4), suggesting a role in oocyte divisions that directly ensue oocyte growth^14,17^. Moreover, the majority of these genes are translated during the first meiotic division^31^ (Table 5). We therefore probed whether compromised splicing in SN oocytes due to disrupted cytoplasmic forces affects the first meiotic division (Fig.4a-b, Extended Data Fig.10a-b). FMN2-mutant oocytes underwent nuclear envelope breakdown comparably to Controls yet predominantly failed to divide, confirming previous observations^32,33^. Mutant cells rescued with FMN2 yet with insufficient time to stir back their cytoplasm before entry into division also predominantly failed to divide. By contrast, restoring cytoplasmic stirring in mutant oocytes for a time sufficient to rescue nuclear droplet and splicing phenotypes led to successful divisions in a majority of cells. We then sought to confirm this implication of splicing, the large majority of which is droplet-associated in SN oocytes, in the success of meiotic division. We treated Control oocytes with Tubercidin, an adenosine analogue previously shown to disrupt nuclear speckles and alter splicing^34^ (Fig.4a-b, Extended Data Fig.10a-j). This treatment gradually decreased droplet size, number, and associated splicing. Within 4 hours, splicing in droplets decreased to FMN2-mutant levels and was altered with comparable tendencies. We then washed-out Tubercidin and monitored the consequences on oocyte division. Resembling FMN2-mutants, the treatments did not affect nuclear envelope breakdown but gradually hindered cell division. Cells that still divided did so with a significant delay, which is a poor prognostic of oocyte fitness. Treating oocytes with Pladienolide B, a potent spliceosome inhibitor specifically targeting the SF3b complex^35^, altered splicing and, after wash-out, also caused cell division delay and failure (Fig4a-b, Extended Data Fig.10a-b and h-j). Thus, cytoplasmic forces in growing oocytes reorganize nuclear droplets across scales to drive subsequent meiotic success critical for fertility.

**Fig. 4.**
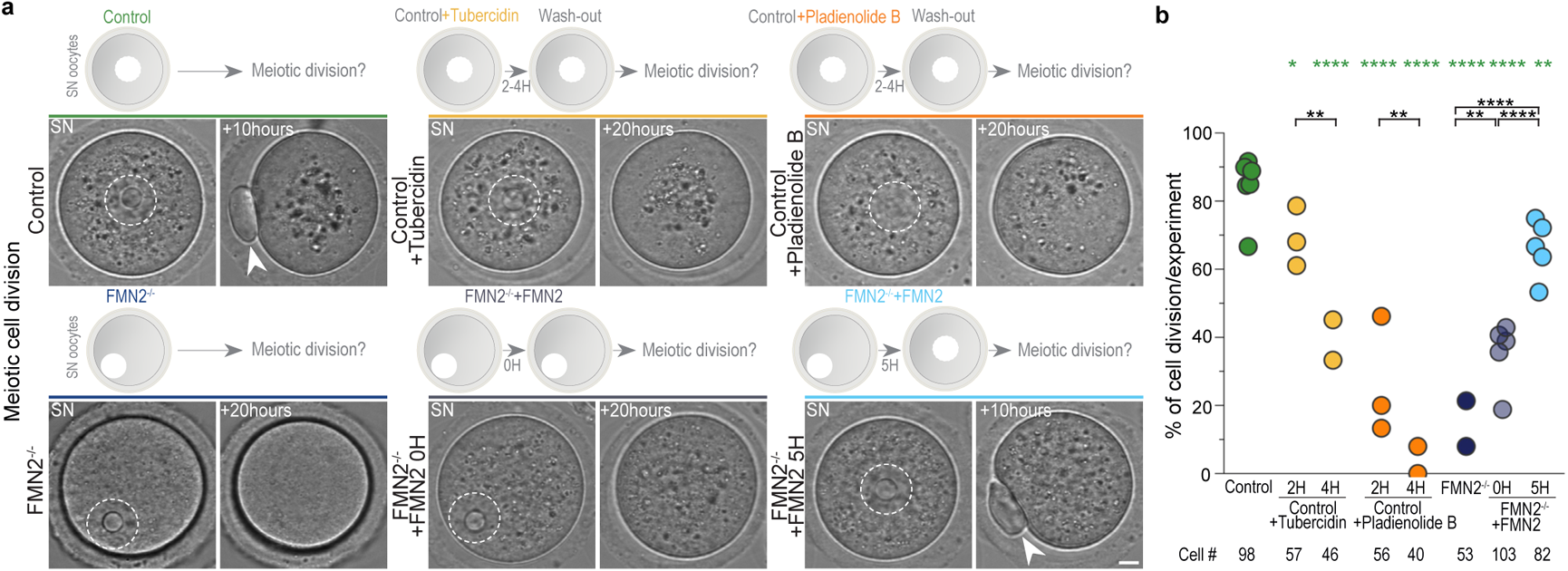
Cytoplasmic force-based regulation of mRNA processing drives meiotic cell division. (**a**) Assessment of the first meiotic division in Control, Control+Tubercidin, Control+Pladienolide B, FMN2^−/−^, FMN2^−/−^+FMN2 0-hour and 5-hour rescue contexts; time-lapse images of Prophase SN stage oocytes 6 minutes prior to entry into division, measured by nuclear envelope breakdown, and 10 to 20 hours after nuclear envelope breakdown with illustrations of each approach above; nuclei are outlined in dashed white; arrowheads indicate the extruded polar body, a readout of successful oocyte division. (**b**) Quantifications of successful oocyte division in Control, Control+Tubercidin, Control+Pladienolide B, FMN2^−/−^, FMN2^−/−^+FMN2 0-hour and 5-hour rescue contexts shown in percentage of divided cells per experiment; cell number depicted below; *P* values derived from Fisher’s exact test of proportions, \**P*=0.0245, \*\**P*<0.0091, \*\*\*\**P*<0.0001. Scale bar, 5µm.

We finally assessed evolutionary conservation of the cytoplasmic capacity to reorganize nuclear liquid compartments. We chose *Drosophila melanogaster* oocytes due to their well described cytoplasmic stirring dynamics^36,37^, yet whose consequences on the nucleus interior remain unexplored. We therefore immunostained nuclear speckles in mid-oogenesis *Drosophila* oocytes (stage 9) when the cytoplasm is randomly stirred by microtubules and buffered by an actin mesh^36,37^, which is the exact opposite of mouse oocytes where microtubules buffer actin-based stirring of the cytoplasm^5,7^. Disrupting the buffering actin mesh precociously via genetic inhibition of two distinct F-actin nucleators Cappuccino or Spire (mouse FMN2 or SPIRE homologues), amplified microtubule-based stirring of the cytoplasm and prematurely induced fast streaming (Fig.5a-b), a known transporting mechanism that, in various animals, plants and fungi, is vital for reproduction or developmental growth and differentiation^36,38^. Premature fast streaming during growth consistently led to larger yet less numerous nuclear condensates by stage 9 when compared to Controls (Fig.5c-e). Inversely, stabilizing the actin mesh by overexpressing a constitutively active Spire blocked cytoplasmic stirring in growing oocytes^36^ that, by stage 9, coherently lacked large nuclear condensates and showed enrichment of smaller condensates (Fig.5a-e). Thus, cytoplasmic actin-based random forces, as in mouse oocytes, or microtubule-based random or streaming forces, as in fly oocytes, can boost nuclear condensate fusion. Moreover, the cytoplasmic capacity to refashion nuclear liquid compartments is evolutionary conserved and could potentially be deployed by other distant species for similar purposes.

**Fig. 5.**
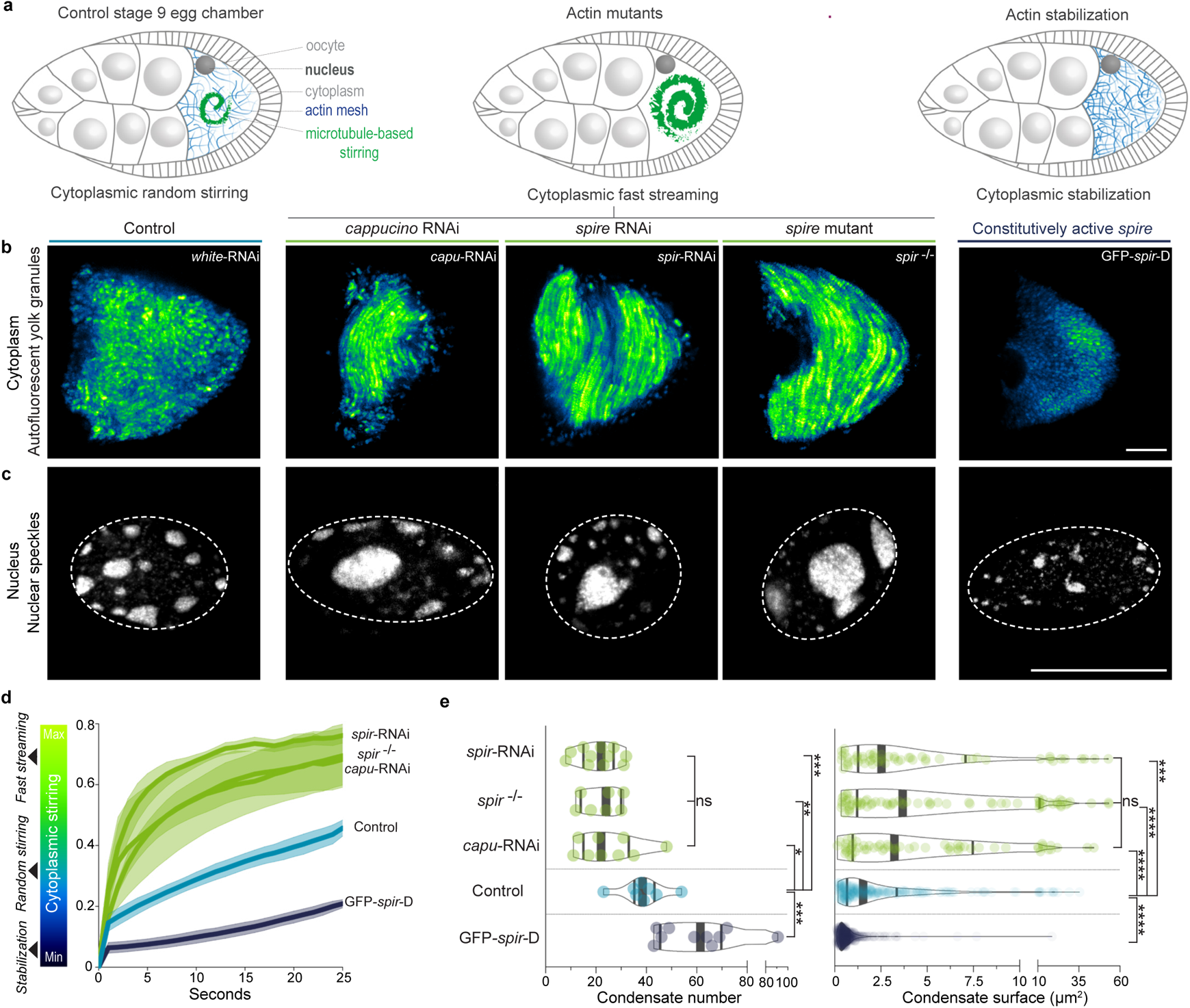
Cytoplasmic aptitude to reorganize nuclear condensates is evolutionary conserved in insects. (**a**) Illustrations of *Drosophila melanogaster* stage 9 egg chambers composed of the oocyte surrounded by nurse and follicular cells. At stage 9, cytoplasmic random stirring occurs in controls (left), premature microtubule-based fast streaming in actin mutants (center), and stabilization in mutants with constitutively active actin nucleators (right). (**b**) Time-projections of cytoplasmic stirring viewed with autofluorescent yolk granules in Control, *cappuccino* and *spire* mutants, or constitutively active *spire* overexpression mutants (GFP-*spir*-D); single z-slice time-projections of 25 seconds shown with ‘GFB’ LUTs. (**c**) Immunoreactivity of speckles in nuclei of oocytes with Control cytoplasmic random stirring, amplified fast streaming, or stabilization; 2µm z-projections with nucleus regions outlined in dashed white. (**d**) Cytoplasmic stirring intensity measured in Control and mutant oocytes by image correlation analyses of cytoplasmic pixel evolution; cell number, Control=4, *cappuccino* RNAi=3, *spire* RNAi=5, *spire*^−/−^=3, GFP-*spir-D*=2; error bars represent mean±s.e.m. (**e**) Quantifications of nuclear speckle number and surface in control and mutant oocytes; cell number, Control=8, *cappuccino* RNAi=9, *spire* RNAi=11, *spire*^−/−^=7, GFP-*spir-D*=10; Condensate number, Control=236, *cappuccino* RNAi=62, *spire* RNAi=55, *spire*^−/−^=74, GFP-*spir-D*=437; violin plots with median±quartiles; *P* values derived from two-tailed Mann-Whitney *U*-Tests or Kruskal-Wallis Tests, ns, not significant, *P*>0.576, \**P*=0.0121, \*\**P*=0.0037, \*\*\**P*<0.001, \*\*\*\**P*<0.0001. Scale bars, 20µm.

This multidisciplinary study uncovers that growing mouse oocytes remodel their cytoplasm to enhance nuclear condensate kinetics concurrently across scales for developmental success (Extended Data Fig.10k). Cytoplasmic stirring forces function simultaneously as nuclear condensate colliders, boosting mesoscale reorganization in the nucleus, and as condensate core particle accelerators, optimizing molecular-scale reactions associated with the condensate’s function like mRNA processing. This nuclear reorganization ensures the success of the ensuing meiotic division, thus conditioning optimal oocyte development for fertilization^14^. This scale-crossing kinetic link between cytoplasmic remodeling and functional nuclear condensate reorganization rationalizes why cytoplasmic remodeling and its consequent nucleus positioning in mammalian oocytes is such a consistent predictor of their embryogenic potential^15,16^. We therefore expect this finding to guide future frameworks of reproductive efforts that include human *in vitro fertilization* and oocyte *somatic cell nucleus transfer*^39^ for agricultural reproductive cloning or endangered species conservation.

Cells can thus physiologically deploy cytoplasmic forces to drive complex reorganization of nuclear condensates and associated optimal biochemical activity tailored to cell fate. This coordinating mechanism couples spatial reorganization of condensates at the mesoscale with functional molecular-scale changes within the same condensates, hence providing a concrete mechanistic example to recent speculations linking different condensate-associated length scales^10^. Although we predominantly focus on how cytoplasmic forces regulate nuclear speckles in oocytes, our study implies the concomitant regulation of several other types of condensates as well as the evolutionary conservation of this mechanism. From an evolutionary advantage standpoint, deploying a single mechanism involving simple physical agitation of the nucleus indeed seems like an optimal solution to the complex problem of simultaneously regulating a variety of nuclear condensates along with their singular biochemical outputs. This mechanism is therefore likely to be relevant for other RNA-processing condensates in the nucleoplasm and other cell types in health or disease^8–13^.

## Acknowledgements

We thank T. Lecuit, V. Lallemand-Breitenbach, A. Meunier, and M. Piel for comments on the manuscript; E. Batsché, N. Beaujean, G. Cecere, H. De Thé, M. Singh, and all members of the Verlhac-Terret laboratory for discussions; E. Anceaume, J. Dumont, P. Mailly, and T. Piolot from the CIRB microscopy platform for imaging support; P. Bassereau for supporting the optical tweezer experiments; J. M. Henderson for advice and assistance with the optical tweezer system and data analyses; the CIRB animal facility for animal care; N. Braure, S. Grosclaude, N.G. Kouakou, and S. Maussion for administrative support; C. Antoniewski, L. Bellenger, N. Naouar, and M. Raymond from the IBPS ARTbio bioinformatics platform for their services; M. Ducom (www.marieducom.com) for designing all illustrations following discussions with A.A.J.. In memory of Rabih Al Jord (1957-2021).

## End Notes

### Funding

The Verlhac-Terret and Huynh laboratories are supported by CNRS, INSERM, College de France and the Bettencourt Schueller Foundation. This work received support under the program « Investissements d’Avenir » launched by the French Government and implemented by the ANR, with the references: ANR-10-LABX-54 MEMO LIFE, ANR-11-IDEX-0001-02 PSL* Research University, ANR-11-LABX0038, ANR-10-IDEX-0001-02 CelTisPhyBio; and through grants from the Fondation pour la Recherche Médicale (FRM label to MHV DEQ201903007796; J.R.H. DEQ20160334884), and the ANR (ANR-18-CE13 to M.H.V.; ANR-16-CE13 to M.E.T.; ANR-15-CE13-0001-01 to J.R.H.). F.C.T. is supported by Institut Curie and CNRS. S.C. received an ARC fellowship. N.S.G. is the incumbent of the Lee and William Abramowitz Professorial Chair of Biophysics. A.A.J. received fellowships from ARC (PDF2017050561) and Labex Memolife 2.0.

### Author contributions

A.A.J. and M.H.V. conceived the project, which was supervised by M.E.T. and M.H.V. A.A.J. designed and performed all experiments on mouse oocytes with assistance from A.E and M.H.V. for microinjections. G.L. performed 3D-computer simulations. F.C.T. performed and analyzed optical trapping experiments. A.E. and C.D.S. performed RT-qPCR experiments. S.C. and J.R.H. did all *Drosophila* oocytes experiments. A.A.J. analyzed all mouse and *Drosophila* experiments. A.A.J. directed the analyses of the RNA-seq data in collaboration with the IBPS ARTbio bioinformatics platform. C.A. performed and analyzed bioinformatical enrichment and spatial correlation analyses. N.S.G. and R.V. contributed with the physical modeling and participated in discussions on data interpretations. A.A.J. wrote the manuscript, which was seen and corrected by all authors.

### Competing interests

The authors declare no competing interests.

### Data & materials availability

All data are in the manuscript and in the Extended Data.

### Code availability

Code description and links are in the Methods.

### Correspondence

To Adel Al Jord

### Extended Data include

Materials and Methods Extended Data Figures 1 to 10

Legends for Extended Data Figures 1 to 10

Legends for Supplementary Videos 1 to 7

Legends for Supplementary Tables 1 to 5

Supplementary references

Separate files: Videos 1 to 7 and Tables 1 to 5

## Extended Data

### Materials and Methods

#### Animals and oocyte collection

##### Mice

All animal studies were performed in accordance with the guidelines of the European Community and were approved by the French Ministry of Agriculture (authorization N°75-1170) and by the Direction Générale de la Recherche et de l’Innovation (DGRI; GMO agreement number DUO-5291). Mice used in this study include: OF1 (Oncins France 1; Charles River Laboratories; 8 to 12 weeks old), C57BL/6 (Charles River Laboratories; 10 to 14 weeks old), and Formin2 (^40^) (FMN2^+/−^ and FMN2^−/−^; 8 to 16 weeks old). FMN2^−/−^ males were crossed with FMN2^+/−^ females to obtain both Control FMN2^+/−^ and Mutant FMN2^−/−^ mice. Ovaries were extracted from mice as previously described^41^ into pre-warmed (37°C) M2+ Bovine Serum Albumine (BSA; A3311, Sigma) medium supplemented with 1µM Milrinone^42^, which maintains growing oocytes arrested in Prophase I. Ovarian follicles were punctured with surgical needles to release growing oocytes from antral follicles (end of oocyte growth^43,44^). Oocytes of different sizes were subsequently collected with a Stripper Micropipette (XLAB Solutions), washed, moved into dishes with fresh medium under mineral oil (M8410, Sigma), mechanically dissociated from follicular cells, and left to stabilize for an hour in the incubator at 37°C before proceeding with experiments. Fully grown oocytes were washed 6 times with and transferred into fresh M2+BSA medium without Milrinone to initiate oocyte *in vitro* maturation (IVM).

##### Drosophila melanogaster

The following stocks were used: v;; UASp–white-shRNA (BDSC #35573), v;; UASp–capu-shRNA (BDSC #32922), v; UASp–spir-shRNA (BDSC #43161), GFP-spirD (BDSC #24767), spir^RP^ cn^1^ bw^1^/CyO (BDSC #5113), b^1^ pr^1^ spir^2F^ cn^1^/CyO (BDSC #8723). BDSC corresponds to Bloomington stock center. Flies were maintained on standard medium in 25°C incubators on a 12 h light/dark cycle. The white-shRNA was used as a control for knock-down experiments since *white* is not expressed during oogenesis. We used nos-GAL4-VP16 driver (BDSC #64188) to perform knock downs or over-expression in the germline. Knock downs were performed at 29°C to increase the efficiency of the GAL4 driver. 3-day-old females were collected and dissected in oil (10S, Voltalef, VWR) for live imaging or in PBS1X for fixed experiments and stage 9 egg chambers were selected based on the morphology of the follicular epithelium.

#### Plasmids, *in vitro* transcription of complementary RNA (cRNA), and oocyte microinjection

Histone 2B-RFP (pRN3-H2B-RFP) was used to visualize chromatin (gift from C. Tsurumi^45^). SRSF2-GFP (a.k.a. SC35; NM_011358) was used to visualize nuclear speckles and was purchased from OriGene Technologies (MG202528). FMN2 (pCS2-FMN2-Myc^5^) was used to rescue cytoplasmic stirring in FMN2^−/−^ oocytes. Plasmids were linearized with appropriate restriction enzymes. The T3 mMessage mMachine (AM1384, ThermoFisher), SP6 mMessage mMachine (AM1360, ThermoFisher), and T7 mMessage mMachine (AM13344, ThermoFisher) transcription kits were used to synthetize capped cRNAs and purified with the RNAeasy kit (Qiagen) as previously described^46^. SRSF2-GFP and FMN2 RNA were polyadenylated using the Poly(A) Tailing kit (AM1350, ThermoFisher). RNA concentrations were measured using a NanoDrop 2000 (ThermoScientific). cRNAs were centrifuged at 4°C for 60 minutes at 13,000 RPM before co-microinjection of 600ng/µl SRSF2-GFP and 125ng/µl H2B-RFP cRNA or 900ng/µl FMN2 into Prophase I oocytes in 37°C M2+BSA+Milrinone medium using an Eppendorf Femtojet microinjector. Oocytes were then incubated for 2 hours for SRSF2-GFP/H2B-RP cRNA translation or 0 to 5 hours for SRSF2-GFP/FMN2 or FMN2 cRNA translation before proceeding with experiments. Exogenous SRSF2-GFP expression profiles were comparable to endogenous nuclear speckles. Note that FMN2 is degraded at meiosis resumption and resynthesized during the first meiotic division^33^. A potential difference of FMN2 amount at the end of meiosis I in the two rescue conditions (0 and 5 hours) prior to meiosis resumption thus cannot explain the extent of each rescue in terms of successful cell divisions. This is rather due to the capacity of FMN2 to rescue cytoplasmic stirring and consequently, cytoplasmic remodeling, nucleus position, nuclear speckle dynamics, and splicing activity before meiosis resumption.

#### Pharmacological inhibitors

The following inhibitors were used: 1,6-Hexanediol^22,47,48^ (Hexanediol; 240117, Sigma-Aldrich) at 1 to 5%, Cytochalasin-D^49^ (CCD; PHZ1063, ThermoFisher) at 1µM, Nocodazole (Noco; M1404, Sigma-Aldrich) at 1µM, Paclitaxel^50^ (Taxol; 580555, EMD Millipore) at 1µM, Pladienolide B^35,51^ at 10µM (CAS 445493-23-2, SC-391691, Santa Cruz Biotechnology), and Tubercidin^34^ at 10µM (TO642, Sigma-Aldrich). Dimethyl sulfoxide (DMSO; D2650, Sigma) was used as a solvent for all inhibitors except Hexanediol before addition into M2+BSA+Milrinone medium. Hexanediol crystals were directly dissolved in the medium at 37°C before incubation with oocytes for 7 minutes (1% for Coilin and TDP-43) or 10 minutes (5% for nuclear speckles). Oocytes were washed 3 times in inhibitor-supplemented medium and incubated for 60 minutes (CCD), 60 to 90 minutes (Noco, Taxol), or 120 to 240 minutes (Tubercidin, Pladienolide B) before proceeding. For droplet coalescence speed experiments, inhibitors were added 10 minutes before start of filming. For oocyte division experiments, oocytes were incubated in inhibitor-supplemented M2+BSA+Milrinone medium for 2 or 4 hours before washing out the inhibitors with M2+BSA medium (without Milrinone) and proceeding with IVM in the incubator or the microscope chamber for up to 20 hours.

#### Immunostainings

##### Mouse oocytes

Prophase I oocytes were incubated 2 to 5 minutes with a 0.4% Pronase (P5147, Sigma) solution in M2+BSA+Milrinone to dissociate the zona pellucida, washed 6 times with fresh M2+BSA+Milrinone medium, and maintained in it for 90 additional minutes. For detection of new transcripts, washed oocytes were incubated with 5-<u>E</u>thynyl <u>U</u>ridine^52^ (EU; 0.5mM) from the Click-IT RNA Alexa Fluor 488 Imaging kit (C10329, ThermoFisher) in M2+BSA+Milrinone for 240 minutes. Before fixation, oocytes were washed three times in M2 medium supplemented with PVP (P0930, Sigma) at 37°C and placed on coverslips coated with gelatin and polylysine. Oocytes were then fixed without permeabilization in paraformaldehyde (PFA 4%; 18814, Polysciences or 15710, Electron Microscopy Sciences) at 30°C for 30 minutes. After a PBS1X wash, oocytes were permeabilized and pre-blocked for 15 min with PBS1X with 0.5% Triton-X (93443, Sigma) and 3% BSA (A2153, Sigma) at room temperature, before incubation with primary antibodies overnight at 4°C followed by an hour incubation at room temperature with secondary antibodies. Primary and secondary antibodies were diluted in PBS1X with 0.2% Triton-X and 3% BSA. Cells were counterstained with DAPI (10 µg/ml; D9542, Sigma) for DNA observation and mounted in Prolong Gold antifade medium (P36941, ThermoFisher) placed in 250 nm thick chambers (70366-12, Electron Microscopy Sciences) adhered to microscope slides (631-1554, VWR) to avoid oocyte squashing. EU revelation was performed before mounting according to the kit instructions. For F-actin visualization, oocytes were fixed as in (^53^) without Pronase treatment, labelled with Phalloidin conjugated with Alexa Fluor-488 (10 U/mL; A12379, ThermoFisher), and mounted in Vectashield Antifade medium (H-1000, Vector Laboratories).

##### Drosophila melanogaster oocytes

Ovaries from 3-day old females were dissected in PBS 1X, fixed in PFA 4% for 20 minutes, permeabilized in PBT (0.2% Triton-X) for 30 min, left overnight with primary antibodies in PBT at 4°C, washed 3 times 30 min in PBT, left with secondary antibody for 2 h at room temperature, washed 3 times 30 min in PBT and mounted in Citifluor medium (Electron Microscopy Sciences) for observation.

##### Antibodies

The following antibodies were used: rabbit anti-Coilin (1:2000; ab210785, Abcam), mouse IgG1 anti-Fibrillarin (1:60; ab11826, Abcam), mouse IgG1 anti-nuclear pore complex proteins MAb414 (1:1000; MMS-120P-100, Eurogentec), rabbit anti-NPAT (1:700; A302-772A, Bethyl Laboratories), mouse IgG1 anti-PSPC1(1:100; clone IL4, SAB4200503, Sigma-Aldrich), rabbit anti-pSF3b155 (phosphorylated at Thr313; 1:400; clone D8D8V, 25009, Cell Signaling), rabbit anti-SF3b155 (1:600; clone D7L5T, 14434, Cell Signaling), mouse IgG1 anti-SMN1 (1:100; Survival of Motor Neurons, clone 2B1, 05-1532, EMD Millipore), mouse IgG1 anti-SRSF2/SC35 (1:400; ab11826, Abcam), mouse IgG1 anti-TDP-43/TARDBP (1:1500; clone 3H8, MABN45, EMD Millipore), and species-specific Alexa Fluor secondary antibodies (1:400; ThermoFisher). A recent study^54^ proposed that the main target of the monoclonal SRSF2/SC35 antibodies is SRRM2 instead of SRSF2/SC35. However, SRRM2 is a spliceosome-associated protein that sharply localizes to nuclear speckles^54^. We also confirmed that the monoclonal ab11826 antibody we used recognized SRS2-GFP^+^ speckles in fixed oocytes expressing SRSF2-GFP. The conclusions of our study are thus unaffected by the proposed SRSF2/SC35 antibody discrepancy^54^.

#### Microscopy

##### Mouse fixed and live imaging

Fixed and live mouse oocytes were examined with a Leica DMI6000B microscope equipped with a Plan-APO 40x/1.25 NA oil immersion objective, a motorized scanning deck and an incubation chamber (37°C), a Retiga 3 CCD camera (QImaging, Burnaby) coupled to a Sutter filter wheel (Roper Scientific), and a Yokogawa CSU-X1-M1 spinning disk. Images were acquired using Metamorph (Universal Imaging, version 7.7.9.0) with 500nm z-steps for a total of 40-50µm in case of fixed cells and 1000nm z-steps for a total of 35-60 µm in case of live cells. Oocytes were placed in a 35mm tissue culture dish with cover glass bottom (FluoroDish FD35-100; World Precision Instruments) for videomicroscopy. Time-lapse 3D-images were acquired at different time intervals (Δt=5 to 12 min) with an exposition time of 500ms for SRSF2-GFP and 300ms for H2B-RFP. For high temporal resolution videos, Metamorph stream acquisition mode was used to capture cytoplasmic random stirring (in bright-field) and rapid diffusive dynamics of nuclear SRSF2-GFP droplets or their surface fluctuations (491nm excitation wavelength) with a Δt of 0.5 seconds on a single z-plane focused on the nucleus or droplet center. For correlations between cytoplasmic stirring intensity and nuclear SRSF2-GFP droplet dynamics, oocytes were first filmed for 120 seconds in bright-field and immediately followed by 120 seconds of filming with a 491nm laser. For IVM experiments, oocytes were filmed in bright-field with a Δt of 3 minutes on a single z-plane focused on the nucleus for up to 20 hours.

##### Optical tweezer setup

The custom-built optical tweezer system was generated by using a near-infrared fiber laser (1064 nm, 5W, Ytterbium fiber laser, IPG Photonics, Oxford, Massachusetts) with a Nikon CFI Plan Apochromat Lambda, 100X, 1.45 NA, oil immersion objective (Tokyo, Japan) mounted on an inverted Nikon C1 Plus confocal microscope (Tokyo, Japan) as previously described^55,56^. The setup is equipped with a temperature controllable stage-top incubator (Tokai Hit STXG-WELSX, Gendoji-cho, Japan) that maintains cells at 37°C in a humidified environment during experiments. Confocal images were acquired using a 488 nm solid-state excitation laser (Coherent, Santa Clara, California) with an ET525/50 bandpass filter (Chroma).

Positions of the trapped cytoplasmic vesicles were detected by recording the light pattern of the outgoing laser in the back focal plane of the condenser with a four-quadrant position-sensitive photodiode (QPD, Hamamatsu, Si PIN photodiode, Product No. S5981). The detected voltage values (V) were converted into displacements and forces by using the conversion value β (µm/V) and the trap stiffness κ (pN/µm)^57^. Ideally, one would determine the conversion value and the trap stiffness for every trapped vesicle. However, the complexity and heterogeneity of the cellular environment, and the occurrence of multiple vesicles being trapped during experiments make it very challenging to do so. We thus determined the value of β by using a polystyrene bead with a diameter of 1 µm to represent the vesicles, by following well-established methods^57^. Briefly, given that the optical trap was stationary, we moved a bead that was stuck to the glass bottom of the cell culture dish across the optical trap laterally by using a piezo stage (Nano-LP100, MadCityLabs, Madison Wisconsin) while recording the output voltage signals by the QPD at 1000 Hz. Within a specific regime where the voltage output is linearly proportional to the stage displacement, we calculated the conversion value β = 1/0.192 µm/V. The trap stiffness was determined by the Boltzmann statistics method^57^. Briefly, we recorded the position distribution of a trapped vesicle by the QPD and then fitted a parabola to the logarithm of the distribution. We obtained trap stiffness κ = 12.8 pN/µm for both x and y directions assuming a symmetric trap. The advantage of the Boltzmann statistic method is that the drag coefficient of the trapped vesicle and the viscosity of the cytoplasm are not required.

To apply forces on the nuclear membranes, we used trapped cytoplasmic vesicles that were located close to the membranes. Given that the optical trap was stationary and the oocytes were immobile, settling on the bottom of the experiment dishes, we moved the piezo stage such that the membranes were moved towards the trapping center while the trapped vesicles were displaced away from the center. Due to the trapping forces of the optical tweezers, the vesicles were then translocated towards the trapping center where membranes were localized, thus pushing against the membranes and consequently exerting forces on the membranes. To calculate the forces (*F*) exerted by the vesicles onto the membranes, we used the displacement of the vesicle according to the equation, *F* = *k* (*r* - *r*_o_), where r_o_ and r are the initial and final vesicle positions, respectively.

##### Fluorescence Recovery After Photobleaching (FRAP)

For laser ablation experiments on SRSF2-GFP droplets, mouse oocytes were imaged at 37°C with a Zeiss (Axio Observer.Z1/7) LSM 980 confocal microscope equipped with Airyscan 2, 2 PhotoMultiplier Tubes (PMT), a GaAsP spectral sensor, a Plan-APO 40x/1.3 oil immersion objective, a 488nm 10mW laser, and a motorized deck with a temperature-controlled chamber. Images were acquired using Zen 3.0 software. A total of 150 frames of 579×579 pixels (53µmx53µm; 16-bit depth) were acquired on single z-planes with bidirectional scan speed of 6 and 1.43 second intervals for a total of 214 seconds. Three frames preceded the bleaching (full 488nm 10mW laser power) of a fixed 8µm×12µm region of SRSF2-GFP droplet-containing nucleoplasm. Bleached region size was fixed to be larger than the target droplet and thus included the neighboring dissolved SRSF2-GFP phase. Fluorescence recovery was imaged with 1.2% laser power.

##### Drosophila melanogaster fixed and live imaging

Drosophila egg chambers were imaged with an inverted spinning-disk confocal microscope (Roper/Nikon) equipped with a sCMOS camera, with a 20X/0.75 objective for live egg chambers, and a 60X/1.4 oil immersion objective for fixed egg chambers. Images were acquired with Metamorph with 1000nm z-steps for a total of 30µm in case of fixed cells. To monitor cytoplasmic stirring, autofluorescent yolk granules in live oocytes were captured on single z-planes every 7 seconds using a 405nm laser.

#### Quantifications and image analyses

Data were obtained from at least 3 independent experiments unless stated otherwise. All images were analyzed on Fiji. All graphs and statistical analyses were generated using GraphPad Prism 9.

##### Cytoplasmic stirring

Cytoplasmic stirring intensity in mouse and *Drosophila* oocytes was determined by image correlation analyses using a previously published software^5^ and available on https://github.com/Carreau/OOCytes/tree/0.9. The software measures pixel changes between consecutive images. Raw time-lapse images (Δt=0.5 seconds for mouse and Δt=7 seconds for *Drosophila* oocytes) were first realigned using the Fiji StackReg plugin. In mouse oocytes, bright-field image correlations were calculated in 3 (NSN) or 4 (SN) cytoplasmic regions of 324µm^2^. In *Drosophila* oocytes, image correlations of autofluorescent yolk granules were calculated in 2 to 5 cytoplasmic regions of 441µm^2^. Correlation values from different regions within a cell were averaged. For visual clarity purposes, final correlation values were transformed by subtracting the value of each timepoint from 1 to obtain an inverted exponential-like curve.

Mouse oocyte cytoplasmic vector maps were generated by the Spatiotemporal Image Correlation Spectroscopy^58^ (STICS) plugin previously implemented for detecting cytoplasmic flows in mouse oocytes^59^ and available on https://research.stowers.org/imagejplugins/. The maps show cytoplasmic flow velocity magnitude and direction. Bright-field time-lapse images (Δt=0.5 seconds) were converted to 32-bit format, realigned using the Fiji StackReg plugin, and masked specifically around the oocyte contour before launching the STICS map jru V2 plugin with box-checking of output velocities, movie mask use, and a time correlation shift of 3 frames.

##### Immunocytochemistry

All immunocytochemistry images spanned 40-50µm (Δz=0.5µm) to include the entire ~30µm wide nucleus and were examined in 3D. Condensate numbers and surfaces were quantified on z-projections covering the entire nucleus for mouse/*Drosophila* nuclear speckles, on z-projections covering the entire condensate signal for Coilin and TDP-43, and on single z-planes for the nucleolus. The maximal radius of nuclear speckles or nucleoli served to calculate their volume. In Control and mutant SN oocytes from the FMN2 mouse strain, 20% to 30% of cells do not present nuclear speckles or any EU incorporation. Total nucleoplasmic signal intensities of condensate markers, corresponding to the sum of condensed and dissolved phases, were quantified on z-projections covering the entire nucleus and normalized by cytoplasmic background signal intensity.

Total nucleoplasmic signal intensities of SF3b155 and pSF3b155 (pThr313) were quantified on z-projections covering the entire nucleus and normalized by cytoplasmic background signal intensity. Total signal intensity of pSF3b155 (pThr313) in nuclear speckle droplets was measured on z-projections covering single droplets, thus integrating the intensity within the droplet volume, and normalized by an equally sized nucleoplasmic signal of pSF3b155 (pThr313) in a region devoid of nuclear speckles and DNA. Nucleoplasmic pSF3b155 (pThr313) was predominantly composed of either droplet-associated or chromatin-associated signal; signal ratios of droplet-associated to chromatin-associated pSF3b155 (pThr313) were obtained from single z-sections. We documented that: cytoplasmic force intensification in oocytes drove an increase in nuclear speckle droplet mobility all over the nucleoplasm in absence of apparent chromatin-association; an important transcriptional drop accompanied chromatin compaction in these final stages of oocyte growth; and that the large majority of splicing activity by the end of growth occurred in nuclear speckle droplets, which are compartments proposed to be sites of spliceosome accumulation for post-transcriptional splicing completion^25,26^. We therefore speculate that, by the end of oocyte growth, splicing activity is predominantly post-transcriptional.

Cytoplasmic actin filament (stained with Phalloidin) density was quantified with the Fiji Tubeness plugin. The plugin calculates a score of how much local pixels represent a tube based on the eigenvalues of the Hessian matrix, with high intensity pixels corresponding to tubular structures and thus, correspond to actin filaments. The plugin also applies a watershed algorithm to measure the average size of the signal void in between the filaments. We verified the expected inverse relation between actin filament density and signal void in our data. Twenty measurements of cytoplasmic regions of 10µmx20µm were made per oocyte on different apico-basal z-planes to maximally cover the entire cytoplasm.

Oocyte volumes were calculated with measured radii values. Nuclear volumes were defined by the external boundaries of the dissolved-phase SRSF2/SC35 signal that occupies the entire nucleoplasm. Briefly, the nucleus was segmented from z-stacks with a Fiji plugin based on signal intensity thresholding of individual z-planes and reconstructed as a 3D-ellipsoid by calculating the minimal-volume enclosing ellipsoid of the thresholded pixels; center position and volume were then calculated from the fitted ellipsoid. The surface occupied by chromatin was quantified on z-projections covering the entire DAPI signal.

In summary, numerous parameters tested in fixed Control and FMN2-mutant oocytes were comparable. This includes: cell growth (*i.e.* cell size, which is also a readout of transcript accumulation in oocytes), nucleus volume, nuclear pore levels (*data not shown*), nuclear g-actin levels^7^, total nuclear protein levels of diverse nuclear condensate markers and the splicing factor CDC5L (*data not shown*), total volume of condensates per nucleus, and EU incorporation as a readout of nascent RNA transcription. This indicates that these parameters are not implicated in splicing changes observed in late growth FMN2^−/−^ oocytes.

##### Droplet tracking

Time-lapse images of oocytes expressing SRSF2-GFP were corrected for bleaching with the histogram matching method and realigned with the Fiji StackReg plugin before proceeding with SRSF2-GFP droplet center tracking using the Fiji Manual Tracking plugin. Temporal Mean Square Displacements (MSD) were calculated from 20 second droplet trajectories. Curves were fitted with the Nelder-Mead method using R software to estimate the diffusion exponent alpha (*α*). As the diffusion was found to be anomalous (*α* <1), we measured the “effective” diffusion coefficient to be able to compare the different conditions. The effective diffusion coefficient *D_eff_* was calculated from a linear fit on the 40 first points (20 seconds) of the temporal MSD curve and normalized by droplet size (*D_eff_* in 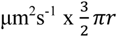 in µm). Although chromatin in fully-grown SN FMN2^−/−^ oocytes is slightly less condensed than in Controls^60^ and may thus interfere with droplet diffusion, Taxol or Nocodazole treatments did not affect the observable chromatin condensation state when compared to Controls thus eliminating the potential chromatin-based bias. In long timescale 3D-tracking, droplet trajectories were corrected for drift with the nucleus centroid trajectories, and displacement distances were squared in graphs. For SRSF2-GFP droplet number and size evolution in NSN oocytes, droplets were manually counted in 3D on every timepoint and maximal droplet surfaces were measured on first and last timepoints. Chromatin (H2B-RFP) surface evolution was quantified in the same NSN cells on 40µm z-projections covering the whole signal and on every timepoint.

##### Droplet surface fluctuations

Droplet contour evolution in time was measured with a custom-built plugin Radioak for use in Fiji and available on https://github.com/gletort/ImageJFiles/tree/master/radioak. The plugin extracted the values of radii of a given selection for all angles around the selection center. Shape variation over time was measured by comparing the value of the radius relative to its average value for each angle. In Fig.3a (upper right), contour fluctuation dynamics were represented visually with a color code: orange if the radius increased or decreased relative to the previous timepoint and white if stable when compared to a given threshold.

In all conditions, SRSF2-GFP droplets of comparable sizes (radius range from 2 to 2.7 µm) were selected. Time-lapse images spanning 15 seconds (Δt=0.5 seconds) were cropped, realigned with the Fiji StackReg plugin, smoothened, and signal thresholded (dark background; default method). The generated binary droplet mask was then analyzed using the Analyze Particles option and the output saved in zip format for Radiak. Radiak was subsequently launched with 1° angle increments from 0 to 360° (input of 360) and the radii values extracted. For each angle, the mean radius “R” over all 30 timepoints was subtracted from the droplet radius “r” for each timepoint. The variance (r-R)^2^ in µm^2^ corresponds to the measure of droplet surface fluctuations that was plotted.

##### FRAP

FRAP image sequences were realigned with Fiji StackReg and SRSF2-GFP droplets within the region of interest were selected for fluorescence recovery analysis. Recovery was measured as fluorescence intensity of photobleached droplet corrected for background and normalized by the droplet’s prebleach intensity. Recovery curves were rescaled from zero (bleach value) to one (100% recovery) and were fit to a simple exponential function (one-phase association) with Prism. Similar to (^61^), the recovery timescale (tau; *τ*) was extracted to calculate the apparent diffusion coefficients (*D_app_*) of droplets with a radius *r* as *D_app_ ~ r^2^/ τ*. Mobile fractions were determined as fluorescence intensity recovery fractions at 120 seconds for NSN and at 60 seconds for SN oocytes, which correspond to Control curve plateaus.

#### Biophysical model

Here, we briefly present a theoretical framework that provides a physical basis for interpreting the observations reported in this study. In (^5^), we described the oocyte cytoplasm as a fluid actively driven out of equilibrium by actomyosin-based mechanical forces. This activity, whereby chemical energy was converted to random active forces, triggered the active diffusion of tracer particles (vesicles) in the cytoplasm. This could be quantified by the mean squared velocity <v^2^> obtained from PIV analyses and was found to be larger than classical thermal diffusion. We argued in (^5,7^) that over diffusion timescales across the oocyte, the cytoplasmic activity could be interpreted as an effective temperature in the cytoplasm defined by T_c_ ∝ <v^2^>. In (^7^), we proposed that cytoplasmic activity enhanced fluctuations of the nuclear membrane which acted as a mechanical transducer, transmitting cytoplasmic active forces to the nuclear interior in the form of effective stirring of the nucleoplasm. This stirring resulted in the active diffusion of chromatin and the chromatin-embedded nucleolus^7^, and could again be interpreted as an effective temperature defined by T_n_ ∝ <v_n_^2^>, where v_n_ is the velocity of intranuclear tracer particles.

Consistent with these earlier findings, we report here that cytoplasmic activity triggered the active diffusion of micron-scale nuclear liquid-like condensates, thereby significantly accelerating their coalescence dynamics. The following scaling argument supports this mechanism. As a readout of the cytoplasmic active forces that are effectively transduced to the nucleoplasm, we use the instantaneous velocity of the nuclear membrane, which can be accessed experimentally: v_nm_~ 0.3 µm/s in SN oocytes. We denote *a* the typical lateral extension of such active deformation of the nuclear membrane, which acts as a localized fluctuating source that stirs the nucleoplasm, and *l*~ 2.7 µm its mean amplitude, measured experimentally in SN oocytes. Assuming in a first approximation a viscous nucleoplasm, a single fluctuating active source at the nuclear membrane acts as a Stokeslet of random direction and intensity and induces a long-range fluctuating flow in the nucleoplasm such that <v_st_^2^*(r)*>~v_nm_^2^ *a^2^/r^2^*, where *r* denotes the distance from the source, and <. > the average over the fluctuating dynamics of the active source. Note that here ~ means proportionality with a dimensionless coefficient of order 1. We next consider that cytoplasmic activity effectively induces a uniform density *c*~1/*a^2^*of independent active sources on the nuclear membrane, which is assumed to be spherical. Here independence of the sources assumes that correlations induced by tension and bending of the nuclear membrane are negligible, which holds for sufficient activity. Summing the contribution of all sources, the fluctuations of the nucleoplasmic flow at a given point at a distance r_0_=uR from the center of the nucleus (of radius R) can be obtained after standard algebra as:

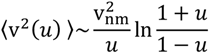

Averaging next over all points in the nucleus shows finally that cytoplasmic activity, transduced by the nuclear membrane, causes a fluctuating flow within the nucleus such that <v_nu_^2^>~v_nm_^2^, where numerical prefactors of order 1 are omitted.

Importantly, the resulting flow spans the entire nucleus (independently of its radius), and has the same order of magnitude as the active sources localized at the nuclear membrane. The impact of such fluctuating flow on the dynamics of nuclear condensates can be quantified by the Peclet number P_n_ =v_nu_ *l*/D_n_, where D_n_ is the mean value of the diffusion coefficient of nuclear condensates. The above orders of magnitude, together with the measured D_n_~ 0.16 µm^2^/s in SN oocytes, yields P_n_~5, which is consistent with an effective diffusion of active origin. Of note, the same analysis applies to molecular transport in the nucleoplasm, taking instead of D_n_ the diffusion coefficient D_m_ of molecular markers (dissolved SRSF2-GFP), which we measured by FRAP in SN oocytes. Similarly, we found D_m_ ~ 0.3 µm^2^/s and thus P_n_~2.5, consistent with an effective diffusion of active origin at the molecular scale in the nucleoplasm. Finally, this shows that cytoplasmic activity can be transduced within the nucleoplasm via a simple, passive (*i.e.* without further energy supply) physical mechanism which enhances the diffusion of nuclear condensates and nucleoplasmic molecules.

Furthermore, we report that the cytoplasmic activity was also transduced down to molecular scales within condensates. Indeed, the liquid-like condensates displayed increased surface fluctuations (consistent with a large effective intranuclear temperature T_n_), as well as a faster turn-over of their molecular content which supposedly facilitated the biochemical processes hosted by the condensates, which in our case correspond to mRNA splicing. The instantaneous velocity of the droplet interface could be estimated from experimental data as v_d_~ 0.9 micron/s in SN oocytes. The fact that v_d_ is of comparable order of magnitude to v_nu_ suggests that droplet surface fluctuations are imposed by shear forces in the nucleoplasm, while capillary forces are negligible. In turn, such surface fluctuations drive flows within droplet condensates; the impact of such flows on molecular dynamics can be, as above, quantified by a Peclet number P_d_ =v_d_ *l_d_* /D_d_, where *l_d_* is the amplitude of fluctuations of the droplet interface and D_d_ the diffusion coefficient inferred from FRAP experiments in SN oocytes. Experimental values yield *l_d_* ~0.5 µm and D_d_~ 0.3 µm^2^/s so that P_d_~1.5, which is consistent with an effective diffusion of active origin at the molecular scale within nuclear liquid-like condensates.

Altogether, our observations suggest an energy cascade triggered by the cytoplasmic actomyosin activity, transduced by the nuclear membrane to the nucleoplasm causing active diffusion and coalescence of liquid-like condensates, and eventually transduced to the liquid-like condensate surface and interior, observed through condensate surface fluctuations and internal molecular turn-over. Based on this physical picture, it can be hypothesized that, due to energy dissipation, the contribution of active forces to fluctuations diminishes along this cascade from the cytoplasm to intranuclear liquid condensate interiors^62^. While we argued above that such energy cascades could in principle occur without extra energy inputs, additional, active mechanisms cannot be ruled out.

#### Computational models and 3D-simulations

##### Agents

To simulate the diffusion of nuclear droplets, we adapted 3D agent-based simulations from our previous work^18^ implemented in C++; also see (^63,64^) for analytical modeling. We chose an agent-based framework for its flexibility, allowing us to test and compare directly different scenario (chromatin presence, chromatin compaction, nucleus shape…). We defined 3 different types of agents: nuclear speckle-like (SRSF2^+^) droplets, a single nucleolus of fixed size, and chromatin-like obstacles. To simplify the model, each agent was represented as a sphere determined by its center position and radius. Mobility analyses of the droplets and the nucleolus in the experiments revealed sub-diffusive motion inside the nucleus and, thus, obstacles were used to simulate molecular crowding in the nucleus. Nuclear obstacle amounts were based on experimental chromatin surface measurements. Potential droplet neo-nucleation and dissolution were excluded from simulations for simplicity. As observed experimentally, droplets could undergo collision-coalescence.

#### Confinement in the nucleus

All agents were confined inside a static spherical boundary representing the nuclear membrane. The confinement inside the nucleus was modelled as a repulsive force effective as soon as the agent was in contact with the membrane (see (^18,65^)).

##### Agent diffusion

Each agent was presumed to diffuse randomly (Brownian motion) and their velocity was dampened by nucleoplasmic friction. Droplet mobility was assumed to obey the Stokes’ law after verifying experimentally that the relation between SRSF2-GFP droplet velocity and their size was coherent with this law. In continuity with our previous work^7^, we modelled the effect of the cytoplasmic stirring activity on the agent diffusion coefficient inside the nucleus as :

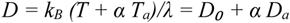

where *D_a_* is the normalized activity-induced diffusion and *D₀* is the diffusion without activity. The values of *D₀* and *α* were estimated by a linear regression on the effective diffusion coefficients of the nucleolus relative to cytoplasmic stirring activity shown in the supplementary data of (^7^).

##### Agent contacts

Contact between agents depended on their type.

- Obstacle or nucleolus with any agent type (droplet, nucleolus, obstacle) : contact between two spheres created a hard-core repulsive force to avoid physical overlap, as in (^18,66^). Repulsion strength increased with sphere overlap, accounting for the limited compressibility of the biological objects^67^.

- Droplet with droplet: droplets coming into close proximity coalesced as soon as they collided, consistent with experimental SRSF2-GFP data. Due to this biological rapidity of SRSF2-GFP droplet coalescence, we rendered coalescence instantaneous in simulations. The coalescing spheres were then replaced by a single sphere with a volume corresponding to the sum of the two original sphere volumes in order to conserve the initial mass.

##### Agent motion

The overall motion of each agent was determined by the balance of all forces it experienced : its intrinsic motility, its contact with other agents, and its contact with the nuclear membrane^18^:

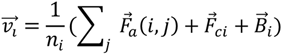

with *η_i_*=6*πriγ* as the friction coefficient opposing agent motion and calculated according to the Stokes’ law, and where *γ* is the viscosity of the medium, *F_a_* is the interaction between agents (hard-core repulsion or attraction), *F_c_* is the force of confinement inside the nucleus, and *B_i_* is the Brownian motion with a diffusion coefficient determined by the cytoplasmic activity.

##### Cytoplasmic activity in Controls±pharmacological inhibitors and mutants

The main effect of the conditions tested in experiments (Control+Nocodazole, Control+Taxol, FMN2^−/−^) was assumed to be the shift in cytoplasmic forces transmitted into the nucleus. In simulations, only the value of the cytoplasmic activity was therefore tuned to mirror these experimental conditions. The values for SN oocytes were all based on experimental data^7^ with Controls being 1, Controls+Nocodazole being 1.9, and FMN2^−/−^ oocytes being 0.2. The values for NSN oocytes were based on the measurements of NSN Control nuclear membrane fluctuations, done as previously^7^ and shown in Extended Data Fig.1j, and with a value estimated at 0.55. The activities of the other biological conditions (inhibitors or mutant) for NSN oocytes were assumed to have a Control-like ratio of activity between SN and NSN oocytes corresponding to a factor of 1.8.

##### Long timescale simulations

In long timescale simulations, we added the possibility for obstacles to adhere compactly to the nucleolus similarly to how chromatin condenses around the nucleolus in physiological conditions of oocyte growth. The obstacles were subsequently constrained to a fixed position relative to the nucleolus whose mobility they follow, and were only permitted small random fluctuations around this position. This allowed us to simulate three different nucleus configurations and test the effect of chromatin compaction on nuclear droplet dynamics. The following simulation series with different nuclear states and a wide range of starting point cytoplasmic stirring activities were performed:

- NSN-like state (simulation series 1): obstacles were widely spread in the nucleoplasm and the starting point intensity of cytoplasmic activity was maintained constant throughout the simulation.

- NSN-like to SN-like transition state (simulation series 2): simulations were first in the NSN-like state with all obstacles widely spread and with low agent mobility. Then at 12 hours, 40 % of obstacles closest to the nucleolus were attracted towards and adhered to the nucleolus, mimicking chromatin condensation that occurs as of the Trans stage into the SN stage. The percentage of obstacles that adhere to the nucleolus was roughly estimated from experimental measurements of the occupied surface of SN chromatin relative to NSN chromatin. Also, cytoplasmic activity at 12 hours was multiplied by 1.8 times to consider the spike of activity measured in experimental conditions during the NSN to SN transition. The transition estimate of 12 hours was defined by the largest number of SRSF2 droplets quantified in the nucleus of a Control Trans-staged oocyte (Extended Data Fig.4a). Due to assimilation of both chromatin condensation and cytoplasmic force intensification, we consider this simulation series with a starting point activity of 0.55 to be the most representative of nuclear droplet kinetics during physiological end of oocyte growth (Control NSN to SN).

- SN-like state (simulation series 3): 40 % of the obstacles adhere to the nucleolus while the remaining 60% remain spread out. Starting point intensity of cytoplasmic activity is maintained constant throughout the simulation.

Droplet dynamics were simulated for 20 hours (simulation series 1 & 3) or for 40 hours (simulation series 2) with longer-term dynamics (up to 100 hours) predicted by a decreasing exponential fit calculated from the last 5 hours of the simulations.

##### FMN2-mutant oocyte specificities of nuclear shape and chromatin compaction

In simulations of the main manuscript, we explicitly model FMN2^−/−^-like cytoplasmic stirring intensity with Control-like nucleus shapes and chromatin compaction to predict the time necessary to reach a 4-droplet state, which corresponds to 59±8 hours. However, fully-grown SN FMN2^−/−^ oocytes also present a less spherical nucleus shape due to microtubules nucleated from the microtubule organizing centers^7^. These SN oocytes also present a ~10% less compact chromatin than Controls^7^ in a nucleus of comparable volume. Chromatin compaction in mutant NSN and Trans oocytes is comparable to Controls. We therefore assessed the effect of these two supplementary differences (nuclear shape and chromatin decompaction) on droplet dynamics in additional 3D-simulations. We found that nuclear shape does not affect droplet dynamics in a context of Control-like cytoplasmic activity, as the time necessary to reach the 4-droplet state remained comparable (Control-like cytoplasmic activity, nucleus shape, and chromatin compaction t=15±8 hours; Control-like cytoplasmic activity and chromatin compaction with FMN2^−/−^-like nucleus shape t=18±6 hours). In contexts of FMN2^−/−^-like cytoplasmic activity, a 10% less condensed chromatin slightly increased the time necessary to reach 4 droplets, and a less spherical nuclear shape increased the time further (FMN2^−/−^-like cytoplasmic activity with Control-like nucleus shape and chromatin compaction t=59±8 hours; FMN2^−/−^-like cytoplasmic activity and chromatin compaction with Control-like nucleus shape t=87±0.5 hours; FMN2^−/−^-like cytoplasmic activity and nucleus shape with Control-like chromatin compaction t>100 hours). Merging all three FMN2^−/−^-like properties together also significantly increased the time necessary to reach the 4-droplet state (FMN2^−/−^-like cytoplasmic activity, nucleus shape, and chromatin compaction t>100 hours). With the additional layers of complexity, these results provide more precise predictions of droplet dynamics in the most “realistic” version of an FMN2-mutant oocyte. Nevertheless, we excluded these data from the computational section of the main manuscript for simplicity and primary focus on the consequences of cytoplasmic forces on nuclear droplet dynamics with Control-like parameters of nucleus shape and chromatin compaction during oocyte growth.

##### Parameter values & sources

| Agent/Parameter | Value | Source/Information |
| --- | --- | --- |
| ***Nucleus*** |  |  |
| Radius | 12 μm | Experimental measurements |
| ***Nucleolus*** |  |  |
| Radius | 4.5 μm ±0.2 | Experimental measurements |
| Friction | 1 | Empirical to obtain diffusion coefficients similar to experimental measurements of nucleoli in Control NSN oocytes |
| Nucleus repulsion | 30 | Strength of confinement; empirical to avoid objects leaving the nucleus while allowing small deformation |
| Sphere repulsion | 30 | Strength of hard-core repulsion with other agents |
| ***Droplet*** |  |  |
| Initial radius | 0.9 $\mu\text{m} \pm 0.01$ | Experimental measurements of nuclear speckles in NSN oocytes |
| Initial number | 55 | Experimental quantifications of nuclear speckles in NSN oocytes |
| Nucleus repulsion | 30 | Strength of confinement; empirical to avoid objects leaving the nucleus while allowing small deformation |
| Sphere repulsion | 10 | Hard-core repulsion with other agents; empirical, with values moderated to allow small deformation |
| Fusion speed | Instantaneous | Simplification based on experimental observations |
| Friction | 0.55 | Empirical to obtain diffusion coefficients similar to experimental measurements of speckles in Control NSN oocytes |
| ***Obstacle*** |  |  |
| Radius | 0.85 $\mu\text{m} \pm 0.4$ | Empirical to confine nucleolus and speckle motion, and to obtain similar diffusion coefficients and anomalous MSD exponents as in NSN experiments |
| Initial number | 875 | Empirical to confine nucleolus and speckle motion, and to obtain similar diffusion coefficients and anomalous MSD exponents as in NSN experiments |
| Nucleus repulsion | 30 | Strength of confinement; empirical to avoid objects leaving the nucleus while allowing small deformation |
| Sphere repulsion | 30 | Hard-core repulsion with other agents; empirical value equal to the nucleolus value |
| Friction | 5 | Empirical to have a lower mobility when compared to other agents |
| Proportion contact | 0 initial<br>0.4 after switch | Evaluated from experimental NSN vs. SN chromatin area in experiments |
| ***Simulation*** |  |  |
| Time step | 0.01 sec | For numerical stability |
| Transition time | 12 h | Time when simulation state switches from NSN-like to SN-like |
| Minimal mobility $D_0$ | 0.01 | Calculated by linear regression of nucleolus diffusion relative to cytoplasmic stirring<sup>7</sup> |
| Coefficient of mobility $\alpha$ | 0.11 | Calculated by linear regression of nucleolus diffusion relative to cytoplasmic stirring<sup>7</sup> |

#### RNA-sequencing, bioinformatics, and RT-qPCR

##### RNA extraction and sequencing

Mouse oocyte RNA extraction and sequencing were performed previously^7^ and can be accessed on the Gene Expression Omnibus (accession number, GEO:GSE103718).

##### Exon usage analysis

For each sample, between 4 and 15 million reads were mapped to the mm10 reference genome using the splice-aware alignment program Hisat2(^68^) (IUC Galaxy wrapper hisat2 v2.1.0). The differential exon usage expression analysis was based on DEXSeq^69^. Briefly, we used the DEXSeq *prepare_annotation.py* script from the Conda environment bioconductor-dexseq==1.28.1 to generate pseudo exons (exon-bins) by *in silico* segmentation of genes using the Ensembl mm10 gene annotation file Mus_musculus.GRCm38.83.chr.gtf. The relative abundance of each exon-bin was calculated with the DEXSeq-Count algorithm by counting all sequenced reads assigned to each exon-bin. Exon-bins related to multiple genes were excluded from further analysis. Finally, the differential usage of exon-bins between FMN2^+/−^ and FMN2^−/−^ conditions was done with DEXSeq (IUC Galaxy wrappers DEXSeq and DEXSeq-Count, v1.24.0.0). Differentially used exon-bins with a Benjamini-Hochberg adjusted p-value (*P_adj_*) threshold of 0.05 were extracted into sheet 1 of Supplementary Table 1 and selected for further RT-qPCR and exon-bin size analyses. Exon bin sizes in sheet 2 of Supplementary Table 1 were plotted relative to their over- or under-representation in FMN2^−/−^ oocytes (*P_adj_*<0.05). Exons from protein coding transcripts for RT-qPCR validation were selected according to high fold changes (>2) and low *P_adj_* (<0.05).

##### Isoform usage analysis

Transcript isoform abundance quantifications were obtained with RSEM^70^, which uses Bowtie2(^71^) for alignment with a custom index built from the mm10 reference genome and the GTF file Mus_musculus.GRCm38.83.chr, and then computes the expression abundance for each isoform (RSEM, ARTbio Galaxy wrapper, Version 0.9.0). Isoform differential expression analysis between FMN2^+/−^ and FMN2^−/−^ conditions was performed for expressed genes (TPM>1) with more than one isoform using the R package IsoformSwitchAnalysisR^72^ (iSAR) which uses counts generated by RSEM and the DEXSeq algorithm to calculate the differential usage of isoforms. Table 2 contains the lists of differentially used isoforms with an FDR-adjusted *P*-value (False Discovery Rate) threshold of 0.05, alternative splicing events per transcript, splicing coordinates, and isoform switch consequences. Differential isoforms generated by exon skipping from a single gene similarly expressed in Controls and Mutants were chosen for RT-qPCR validation of alternative splicing. Using this approach, we detected 1259 transcripts with altered spicing patterns in FMN2^−/−^ oocytes (Table 2), which probably start appearing by the end of growth as of the Trans-stage when splicing changes become visible (Fig.3d-e). This number of transcripts is coherent with studies reporting ~ 7000 to 11000 expressed genes during the whole process of mouse oocyte growth^73,74^.

##### Enrichment and spatial correlations of SRSF1 and SRSF2 binding sites

SRSF1 and SRSF2 binding sites were obtained from (^28^) for binding site enrichment analyses. The numbers of SRSF1 and SRSF2 binding sites in differentially used exons (DEXSeq) or transcripts (iSAR) were then compared to the number of SRSF1 and SRSF2 binding sites in all genes. Correlation tests of distance metrics between genomic features were implemented using the GenometriCorr package^75^ which computes spatial association of datasets to highlight potentially relevant relationships between them. We compared sites of differential exon usage (DEXSeq) or alternative splice sites (iSAR) with binding sites of SRSF1, SRSF2, MBNL3, and YY1(^28–30^). Two *in silico* controls were also generated (Prom50 and Term50) which correspond to the first and last 50 nucleotides of all RefSeqNCBI transcripts, respectively. When necessary, datasets were converted to the Ensembl mm10 gene annotation using the IUC Galaxy wrapper crossmap.bed v0.5.2+galaxy0 (http://crossmap.sourceforge.net).

##### Gene ontology and analyses of translational status of transcripts

The list of genes affected by differential exon (DEXSeq list) or isoform usage (iSAR list) were merged for functional enrichment analyses and for comparative analyses with transcripts translated during the first meiotic division (Supplementary Tables 4 and 5). Biological processes and Gene Ontologies were analyzed using Enrichr^76,77^ available through the web interface https://maayanlab.cloud/Enrichr/. The full lists of translated, activated (engaged in translation), and repressed (degraded) transcripts were kindly shared by Marco Conti and based on their previous work^31^. Their probe set GeneID’s were updated and converted using the Mouse Genome Informatics website (http://www.informatics.jax.org/batch) before comparison with the FMN2^−/−^ list of genes shown in Supplementary Table 5.

##### RT-qPCR

Total cellular RNA (25 oocytes per sample) was extracted with the RNAqueous-Micro Total RNA Isolation Kit (AM1931; ThermoFisher) following the manufacturer’s protocol and eluted into 20µl of elution buffer before DNAase I treatment. cDNA was synthesized using the iScript Reverse Transcription Supermix (1708840; Bio-Rad) and the quantitative PCR was performed in triplicate with primer pairs (listed below) in a CFX96 Touch Real Time PCR Detection System (Bio-Rad) followed by analysis on the CFX Maestro Software (Bio-Rad). Samples were normalized to that of Rpl19 (Ribosomal protein L19) and Gapdh (Glyceraldehyde 3-phosphate dehydrogenase), and relative fold changes were calculated using the 2^−ΔΔC^ method. For DEXSeq exon usage validation with RT-qPCR, we designed the two primers against the same exon detected as differentially used by DEXSeq. The expression of total transcript was verified (except for Kctd20) with primer pairs against junctions of constitutive exons, which were not classified as differentially used by DEXSeq. The general primer design strategy is illustrated in sheet 2 of Supplementary Table 1. For iSAR isoform usage validation with RT-qPCR, we designed isoform specific primer pairs against exon 2-3 & exon 4 for the long isoform and exon 2 & exon 3-6 for the short isoform, as illustrated in sheet 5 of Supplementary Table 2.

The following primers were used:

Ercc5 (ENSMUSE00000812268) forward primer (5’to3’) F=TCTAAGGAGAGGAACTCAGGGG, reverse primer (5’to3’) R=TCTGCTAGATCATCACTGCTGC; Ercc5 (NM_011729.2) F=GCGTCCTTTATCCTAACGGGA, R=GCCAAATGCTAATATCCACGGC; Hnrnpdl (ENSMUSE00000825422) F=TCTATCTCTGGGGGTCGCAC, R= CTTTACGCTGGTACATGAAGTTGG; Hnrnpdl (XM_036165266.1) F=ATAGGTTCTGGGAAGTGCGA, R=TTGGTTCCAGTTTTGGCCCT; Kctd20 (ENSMUSE00000788494) F=AGATCAAGAGGAGACCTGGCG, R=ACATGGCGACTCTTTCCTTCC; Ncapg2 (ENSMUSE00000266378) F= TCATCCATGTCATCCGCCAC, R= TGGCATTCTCCTTCTCGCATT; Ncapg2 (NM_133762.4) F= GGACCTGATGCAGACTACGG, R=AGGGAGCCTTACAACCCCAG; Ogdh (ENSMUSE00001059524) F=TTAAGGCCATTGACAGCCTCC, R=TACAGGTGCAGAATAGCACCG; Ogdh (NM_001252283.1) F=CCCCTTTCCCTGAGTCGAAG, R=TGGTGACCCCTGACCTGATA; Exosc1 isoform 1 (NM_025644.4) F=AGAATGGCGCGGTTCCC, R=GGCAAACCGTGAGTTGATGC; Exosc1 isoform 2 (NM_001164561.1) F=TGAAGACCAGCGAGAATGGC, R=AATTTCTACCTTACAGGTGACGAC.

## Extended Data Figures 1→10 & associated figure legends

**Extended Data Fig. 1.**
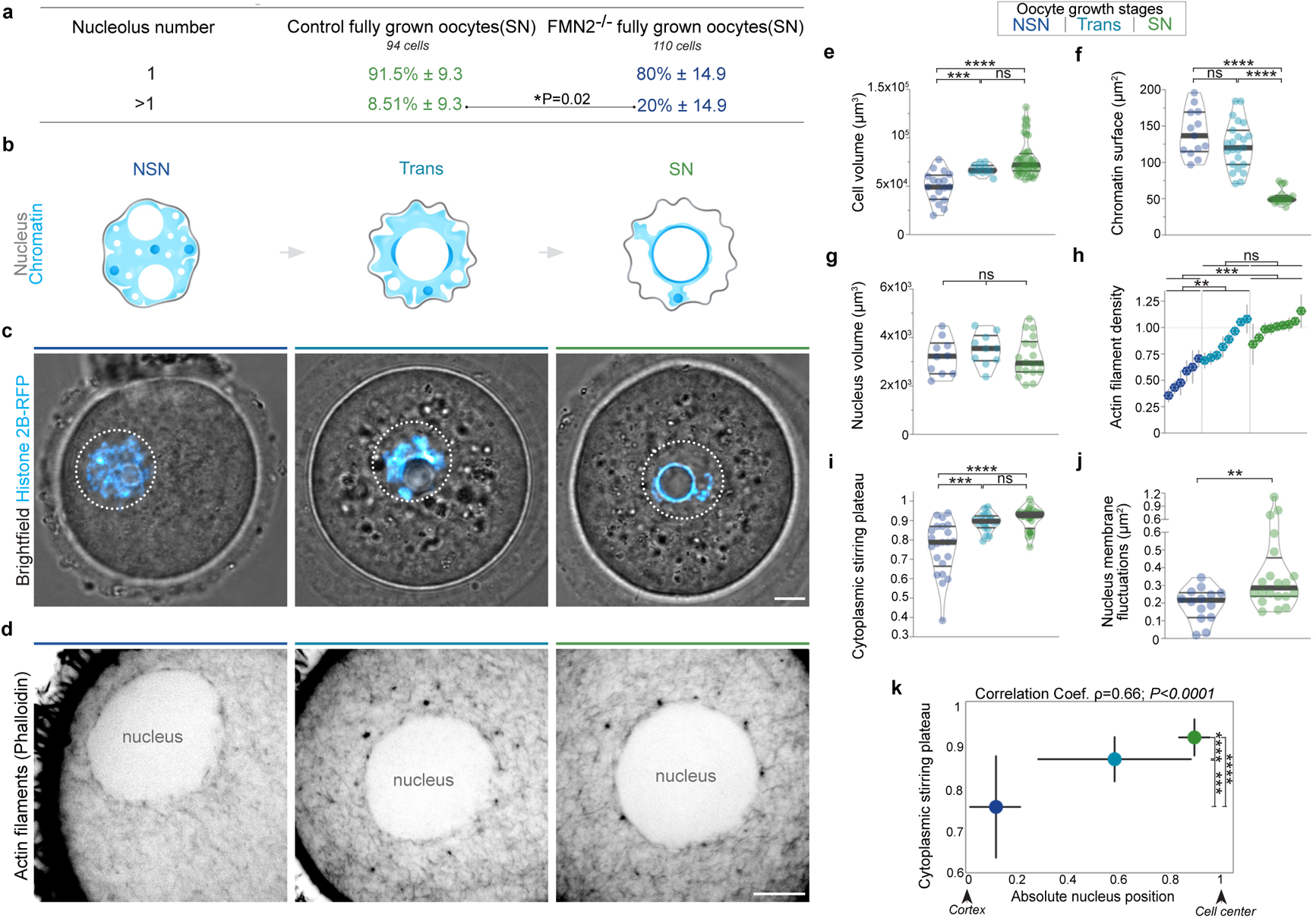
Nucleolus number in Control and F-actin mutant fully grown mouse oocytes and cytoplasmic random stirring relative to oocyte growth stages. (**a**) Nucleolus number quantification showing the ~2.35-fold enrichment in the proportion of cells presenting more than one nucleolus in fully-grown SN FMN2^−/−^ oocytes when compared to SN Controls. (**b**) Illustration of chromatin configuration evolution (NSN-Trans-SN see (^20,78,79^)) coinciding with oocyte growth subcategories (mid-antral, late-antral, and fully grown). The nuclear membrane (gray) is depicted according to its fluctuation intensity at distinct stages of growth (also see (^7^) and (j)). (**c**) Merge of bright-field and fluorescent labeling of live control oocytes expressing the chromatin marker H2B-RFP at distinct growth stages (0.5µm z-planes), linking chromatin configuration and nucleus position evolution with mouse oocyte growth progression; nucleus region is outlined in white. (**d**) Actin filaments (F-actin) stained with Phalloidin relative to oocyte growth progression; note the increase of cytoplasmic F-actin network density as of the Trans stage coinciding with nucleus centering; F-actin is absent from the nucleus interior, similar to what can be observed in human oocytes^80^. (**e**) Quantifications of cell volume in mid-antral (NSN), late-antral (Trans), and fully grown (SN) fixed oocytes; cell number, NSN=17, Trans=11, SN=43. (**f**) Chromatin surface relative to oocyte growth; measurements were made on 40µm z-projections of fixed cells; cell number, NSN=13, Trans=23, SN=18. (**g**) Nucleus volume quantifications relative to growth stages in fixed oocytes; cell number, NSN=10, Trans=10, SN=18. (**h**) Quantifications of cytoplasmic F-actin density per cell relative to oocyte growth stages; 20 measurements of Phalloidin-stained F-actin per cell represented by mean±s.d. spanning different z-planes from top to bottom of each cell and values normalized to the mean SN density value; distribution medians used to compare stages statistically. (**i**) Cytoplasmic stirring intensity plateaus relative to growth stage obtained with image correlation analyses of cytoplasmic stirring in live oocytes; cell number, Control NSN=22, Trans=19, SN=28. (**j**) Nucleus membrane fluctuations in live NSN and SN oocytes measured as in (^7^); cell number, NSN=14, SN=20. (**k**) Cytoplasmic stirring intensity plateaus correlated to absolute nucleus positions in live growing oocytes; absolute nucleus position corresponds to 1-*ρ,* with *ρ* quantified as in (^15^); cell number, NSN=16, Trans=16, SN=20; error represents mean±s.d., Spearman correlation coefficient ρ=0.66, \*\*\*\**P*<0.0001. Violin plots with median±quartiles; *P* values derived from Fisher’s exact test of proportions (**a**), two-tailed Mann-Whitney *U*-Tests (**e, f, g, i, j**, and **k** for nucleus position tests), and Kruskal-Wallis Test (**h**), ns, not significant, *P*>0.0735, \*\**P*<0.0068, \*\*\**P*<0.0006, \*\*\*\**P*<0.0001. Oocyte growth color codes based on cytoplasmic stirring intensities; scale bars, 5µm.

**Extended Data Fig. 2.**
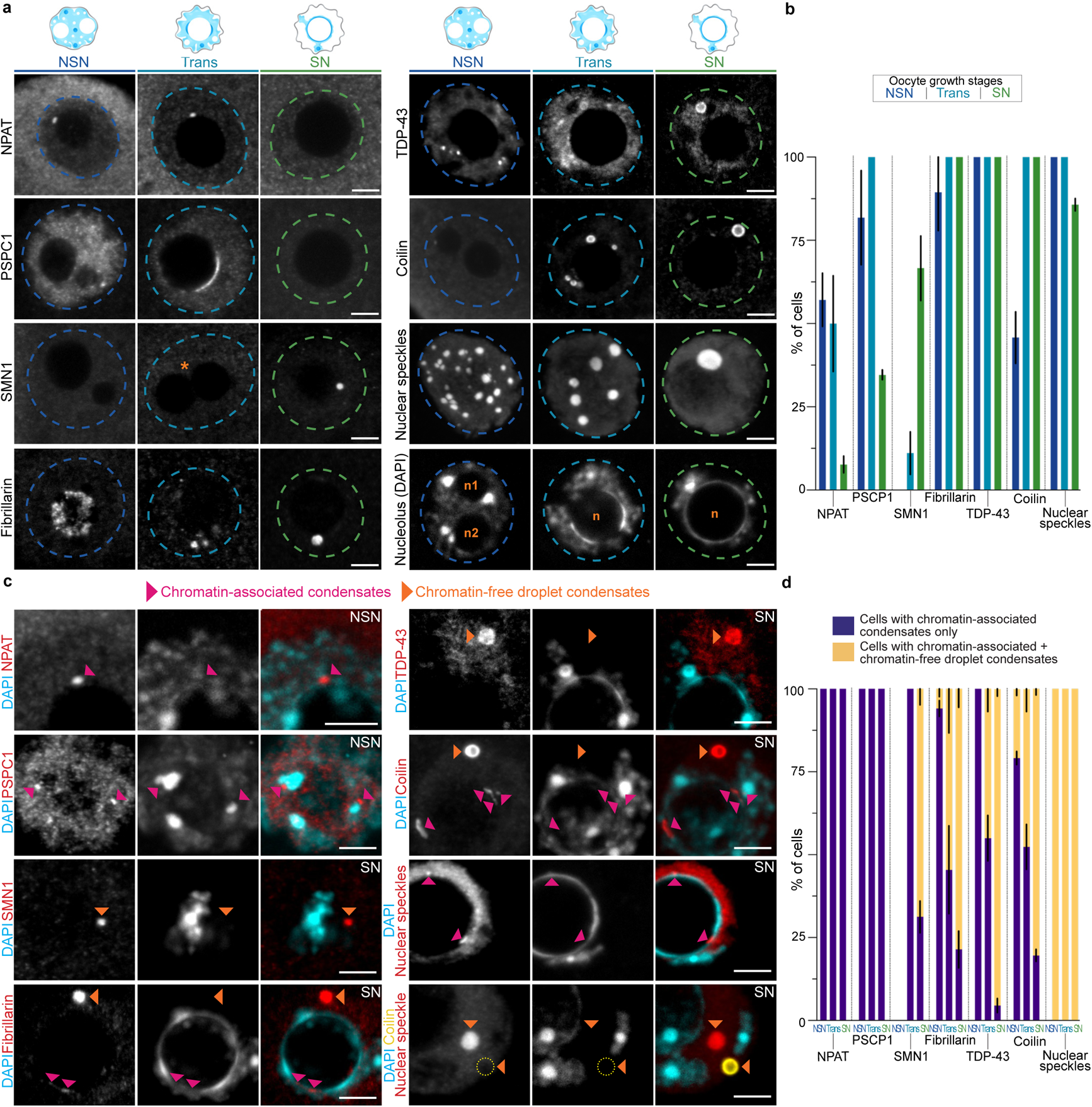
Immunocytochemistry screen of diverse nuclear biomolecular condensates in growing oocytes with classification into chromatin-associated and chromatin-free (droplet) condensates. (**a**) Representative immunostainings of screened nuclear biomolecular condensates using known immunomarkers^21,81^ relative to oocyte growth stages; NPAT (Nuclear Protein, Coactivator of Histone Transcription) is a marker of Histone Locus Bodies, PSPC1 (Paraspeckle Component 1) of paraspeckles, SMN1 (Survival of Motor Neuron 1) of gems and rarely associated with Cajal bodies, Fibrillarin of nucleoli and rarely Cajal bodies, TDP-43 (TARDBP; TAR DNA-Binding Protein 43) of paraspeckles, of TDP-43 bodies and of Cajal bodies, Coilin of Cajal bodies, and nuclear speckles stained with Abcam ab11826 antibody (SC35/SRSF2); Fibrillarin is notably nucleolar in NSN cells before becoming mostly nucleoplasmic as of the Trans stage suggesting the restructuring of the multiphase nucleolus^82^; due to this nucleolar phase restructuring observed as of the Trans stage, the nucleolus (marked with the letter “n”) loses Fibrillarin positivity and becomes recognizable only by signal exclusion and the surrounding DNA (stained with DAPI); asterisk in the SMN1 row indicates a visible nucleolar fusion; single 0.5µm z-planes are shown except for nuclear speckles which are 20µm z-projections; nucleus regions are outlined with dashed lines color coded relative to cytoplasmic stirring intensities at different growth stages. (**b**) Proportions of NSN, Trans, and SN oocytes that contain tested nuclear biomolecular condensates in their nucleus; cell number, NPAT NSN=14, Trans=6, SN=13, PSPC1 NSN=11, Trans=8, SN=26, SMN1 NSN=16, Trans=9, SN=24, Fibrillarin NSN=19, Trans=11, SN=14, TDP-43 NSN=9, Trans=10, SN=22, Coilin NSN=24, Trans=22, SN=56, nuclear speckles NSN=37, Trans=24, SN=42. (**c**) Images from the immunocytochemistry screen showing the two subpopulations of nuclear biomolecular condensates observed at various stages of oocyte growth; smaller, irregularly shaped chromatin-associated condensates indicated by fuchsia arrowheads and larger, round chromatin-free droplet condensates indicated by orange arrowheads; DNA is stained with DAPI; all images are z-projections, 3µm for NPAT and PSPC1, 2µm for SMN1, 5µm for Fibrillarin and bottom SRSF2, 10µm for TDP-43 and Coilin, and 1µm for top nuclear speckles; note that in the bottom nuclear speckles merge, a Coilin droplet is shown in yellow; note also that the nucleolus is an exception to the binary rule of classification since it always is a droplet that is in tight contact with and surrounded by chromatin. (**d**) Proportions of NSN, Trans, and SN oocytes containing nuclear condensates that are exclusively chromatin-associated (purple) or containing both chromatin-associated condensates and droplets (yellow); tested condensate immunomarkers and growth stages are indicated below the histogram; cell number, NPAT NSN=8, Trans=3, SN=1, PSPC1 NSN=9, Trans=8, SN=9, SMN1 NSN=0, Trans=1, SN=16, Fibrillarin NSN=17, Trans=11, SN=14, TDP-43 NSN=5, Trans=20, SN=22, Coilin NSN=11, Trans=21, SN=56, nuclear speckles NSN=37, Trans=24, SN=42. Bars above images are color coded relative to cytoplasmic stirring intensities at corresponding growth stages; scale bars, 5µm.

**Extended Data Fig. 3.**
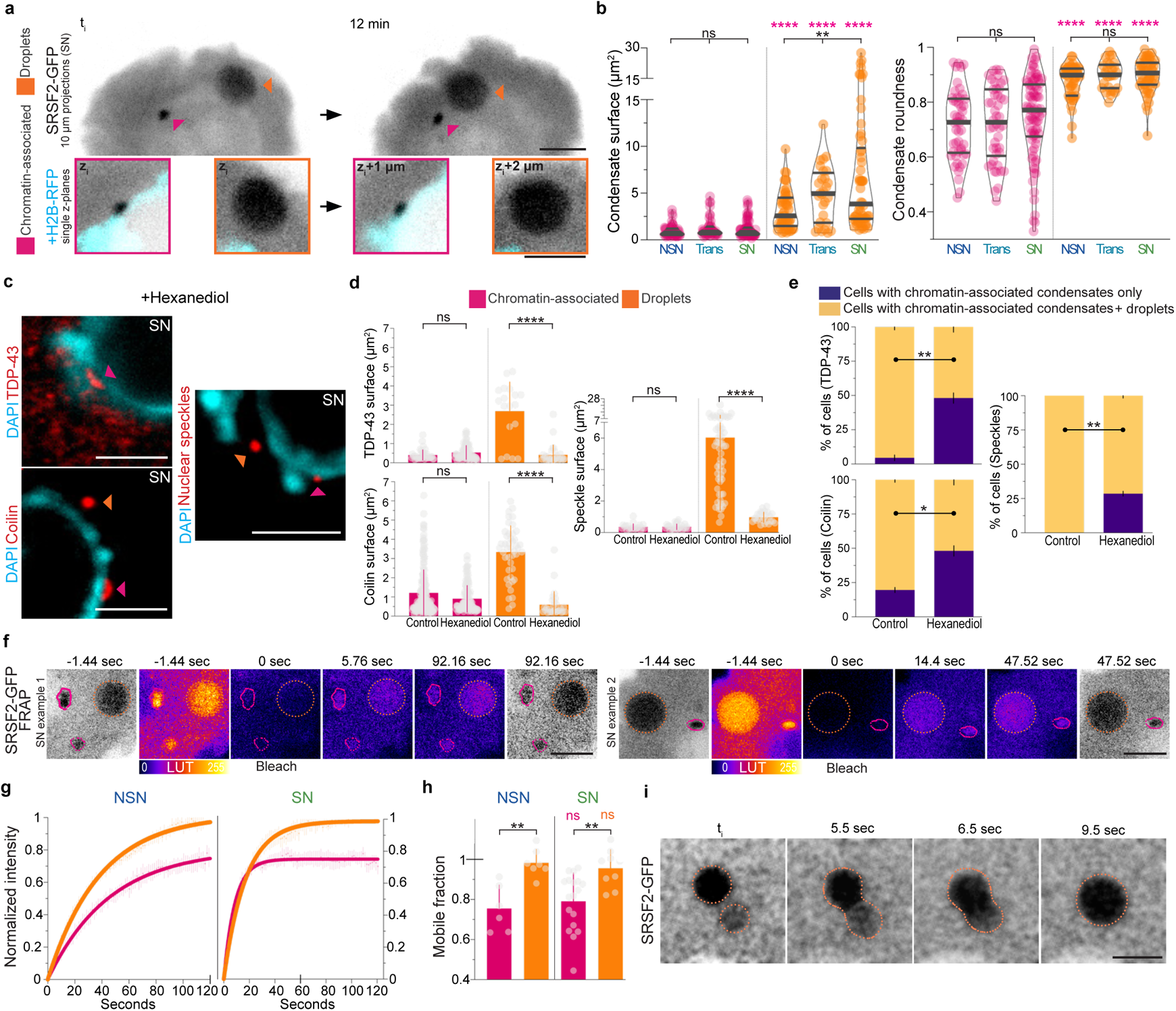
Droplet condensates behave like liquids whereas chromatin-associated condensates do not. (**a-b**) Condensate subtype mobility, size and shape. (**a**) Time-lapse frames showing high spatiotemporal mobility of a large SRSF2-GFP droplet (orange) and low mobility of a small chromatin-associated condensate (fuchsia) that typically remains stably bound to chromatin (H2B-RFP) in a live oocyte. (**b**) Quantifications of surface and roundness of SRSF2-GFP chromatin-associated or droplet condensates in live oocytes at all stages of growth. Droplets are significantly bigger and rounder at all stages of growth; condensate number, chromatin-bound NSN=49, Trans=46, SN=82, droplets NSN=49, Trans=28, SN=58. (**c-e**) Treatment with 1-6-hexanediol, a disruptor of weak hydrophobic molecular interactions that are key for liquid-like behavior^22,47,48^, dissolves droplets without affecting chromatin-bound condensates. (**c**) Representative immunostainings of TDP-43 (0.5µm z-plane) and Coilin (5µm z-projection) in SN oocytes treated with 1-6-hexanediol [1%] for 7 minutes (TDP-43 and Coilin) or [5%] for 10 minutes (nuclear speckles), and showing Control-like chromatin-associated condensates (fuchsia arrowheads) and small droplets (orange arrowheads); DNA stained with DAPI (in blue). (**d**) Quantifications of TDP-43,Colin, and nuclear speckle condensate surfaces in SN oocytes following 1-6-hexanediol [1% or 5%] treatments and showing a size-decrease only for droplets; condensate number, TDP-43 chromatin-associated|droplets, Control=38|18, Hexanediol=84|29, Coilin chromatin-associated|droplets, Control=183|44, Hexanediol=108|29, nuclear speckles chromatin-associated|droplets, Control=22|62, Hexanediol=26|25. (**e**) Proportion of SN oocytes containing TDP-43 and Coilin condensates that are exclusively chromatin-associated (purple) or containing both chromatin-associated condensates and droplets (yellow); cell number, TDP-43 Control=22, Hexanediol=27, Coilin Control=56, Hexanediol=27, nuclear speckles Control=42, Hexanediol=14. (**f-h**) High fluorescence recovery of SRSF2-GFP droplets in NSN and SN oocytes. (**f**) Representative FRAP sequences of SRSF2-GFP chromatin-associated condensates (fuchsia) and droplets (orange). (**g**) Normalized fluorescence intensity recovery curves (mean±s.e.m.) with simple exponential fits of SRSF2-GFP chromatin-associated condensates and droplets in (left) NSN or (right) SN oocytes. (**h**) Mobile fractions of total populations for SRSF2-GFP relative to condensate sub-type and oocyte growth stage; mobile fractions correspond to recovery fractions evaluated at 120 seconds for NSN condensates and 60 seconds for SN condensates; condensate number, chromatin-associated NSN=6, SN=18, droplets NSN=7, SN=9. (**i**) Time-lapse images showing rapid SRSF2-GFP droplet coalescence in a live SN oocyte. Violin plots with median±quartiles (**b**), error bars represent mean±s.d. (**d** and **h**) and mean±s.e.m. (**e**); *P* values derived from two-tailed Mann-Whitney *U*-Tests (**b**, **d**, and **h**), Kruskal-Wallis Tests (**b**), or Fisher’s exact test of proportions (**e**), ns, not significant, *P*>0.0648, \**P*=0.0102, \*\**P*<0.0054, \*\*\*\**P*<0.0001; scale bars, 5µm.

**Extended Data Fig. 4.**
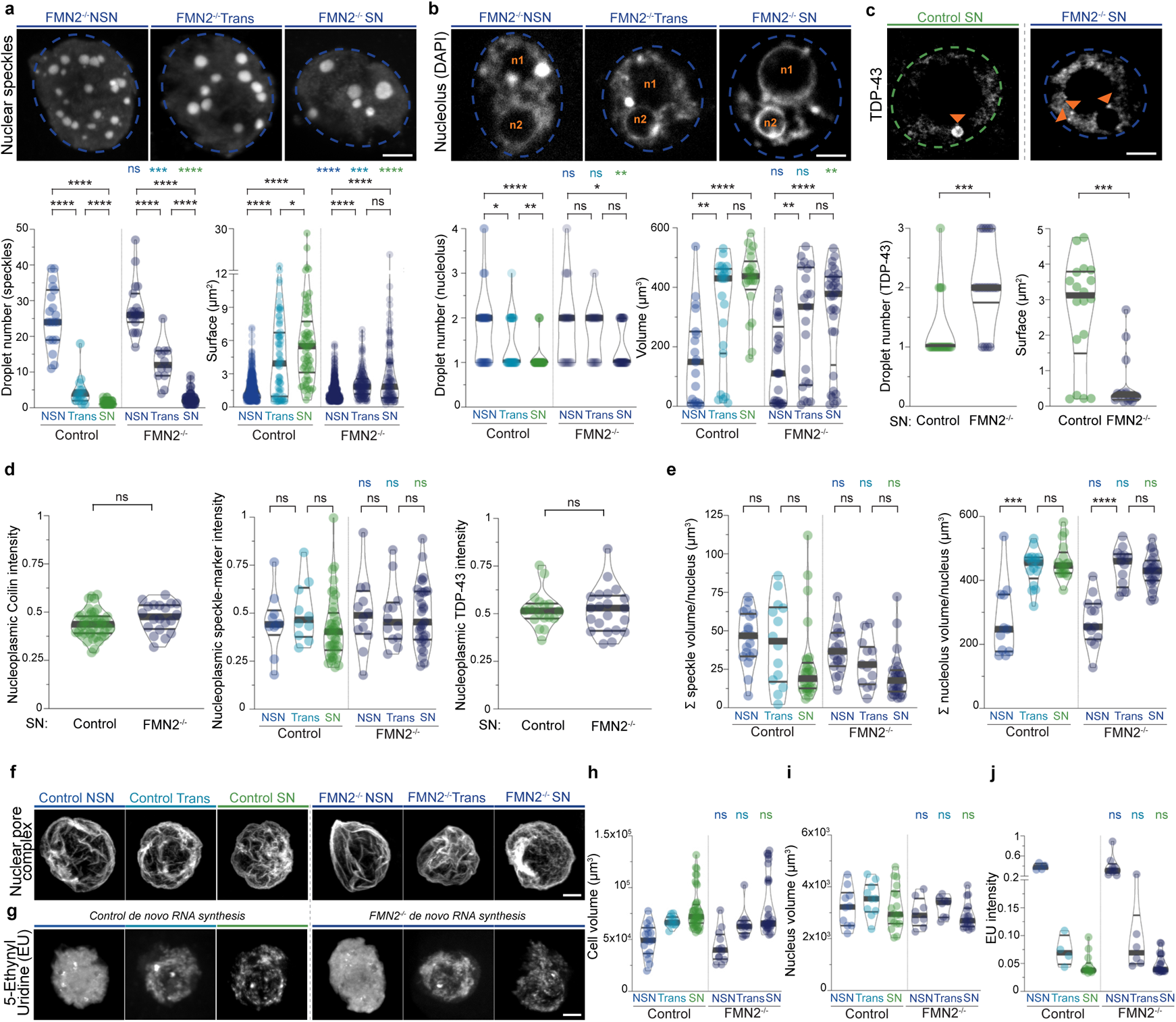
Nuclear droplets are smaller and more abundant in growing F-actin mutant oocytes with disrupted cytoplasmic forces, but total condensate marker amounts, cell or nucleus volumes, and neo-transcription are comparable to Controls. (**a**) Immunostainings of nuclear speckles in growing F-actin mutant (FMN2^−/−^) oocytes and quantifications of droplet numbers and sizes in Control and FMN2^−/−^ NSN, Trans, and SN oocytes (related to Fig.1d-e, also see Extended Data Fig.2a for Controls); cell number (left graph), Control NSN=19, Trans=15, SN=41, FMN2^−/−^ NSN=19, Trans=12, SN=52; condensate number (right graph), Control NSN=450, Trans=69, SN=62, FMN2^−/−^ NSN=528, Trans=149, SN=122. (**b**) Images showing nucleoli and DNA (DAPI) in growing FMN2^−/−^ oocytes; nucleoli visible by signal exclusion and are marked with the letter “n” (see Extended Data FigED. 1a and 2a for Controls); graphs are quantifications of nucleolus numbers and volumes; cell number (left graph), Control NSN=38, Trans=42, SN=66; FMN2^−/−^ NSN=19, Trans=17, SN=36; nucleoli number (right graph), Control NSN=19, Trans=23, SN=21, FMN2^−/−^ NSN=25, Trans=21, SN=32. (**c**) Immunostainings of TDP-43 and quantifications of droplet numbers and sizes in SN Control and FMN2^−/−^ oocytes; orange arrowheads indicate droplets; cell number (left graph), Control=20, FMN2^−/−^=18, condensate number (right graph), Control=18, FMN2^−/−^=15. (**d**) Total nucleoplasmic intensities for Coilin (left), nuclear speckles (center) and TDP-43 (right) in Control and FMN2^−/−^ SN oocytes; cell number for Coilin, Control=43, FMN2^−/−^=20; cell number for nuclear speckles, Control NSN=10, Trans=10, SN=40, FMN2^−/−^ NSN=12, Trans=12, SN=31; cell number for TDP-43, Control=18, FMN2^−/−^=21.(**e**) Sum of nuclear speckle (left) or nucleolus (right) volumes per nucleus in Control and FMN2^−/−^ SN oocytes; cell number for nuclear speckles, Control NSN=19, Trans=14, SN=29, FMN2^−/−^ NSN=19, Trans=13, SN=27; Nucleolus, Control NSN=11, Trans=17, SN=17, FMN2^−/−^ NSN=13, Trans=13, SN=23. (**f**) Representative images of the nucleus membrane immunostained with the Mab414 antibody recognizing the nuclear pore complex in growing Control or FMN2^−/−^ oocytes (top; 20-24µm z-projections). (**g**) Incorporation profiles of 5-<u>E</u>thynyl <u>U</u>ridine^52^ (<u>EU</u>, 1mM incubated for 240 minutes) that reflect *de novo* RNA synthesis in growing Control or FMN2^−/−^ oocytes; 20µm z-projections show comparable drops in transcription with growth progression. (**h-j**) Quantifications of cell volume (**h**), nucleus volume (**i**), and EU signal intensity (**j**) in growing Control or FMN2^−/−^ oocytes; cell number, (**h**) Controls same as in Extended Data Fig.1e, FMN2^−/−^ NSN=12, Trans=12, SN=31, (**i**) Controls same as in Extended Data Fig.1g, FMN2^−/−^ NSN=8, Trans=8, SN=17, (**j**) Controls NSN=4, Trans=4, SN=14, FMN2^−/−^ NSN=9, Trans=16, SN=20. Nucleus regions in a-c outlined with dashed lines; violin plots with median±quartiles; *P* values derived from two-tailed Mann-Whitney *U*-Tests, ns, not significant, *P*>0.0559, \**P*<0.0319, \*\**P*<0.0097, \*\*\**P*<0.0007, \*\*\*\**P*<0.0001; colored “ns” and asterisks are statistical comparisons with Controls; oocyte growth color codes based on cytoplasmic stirring intensities; scale bars, 5µm.

**Extended Data Fig. 5.**
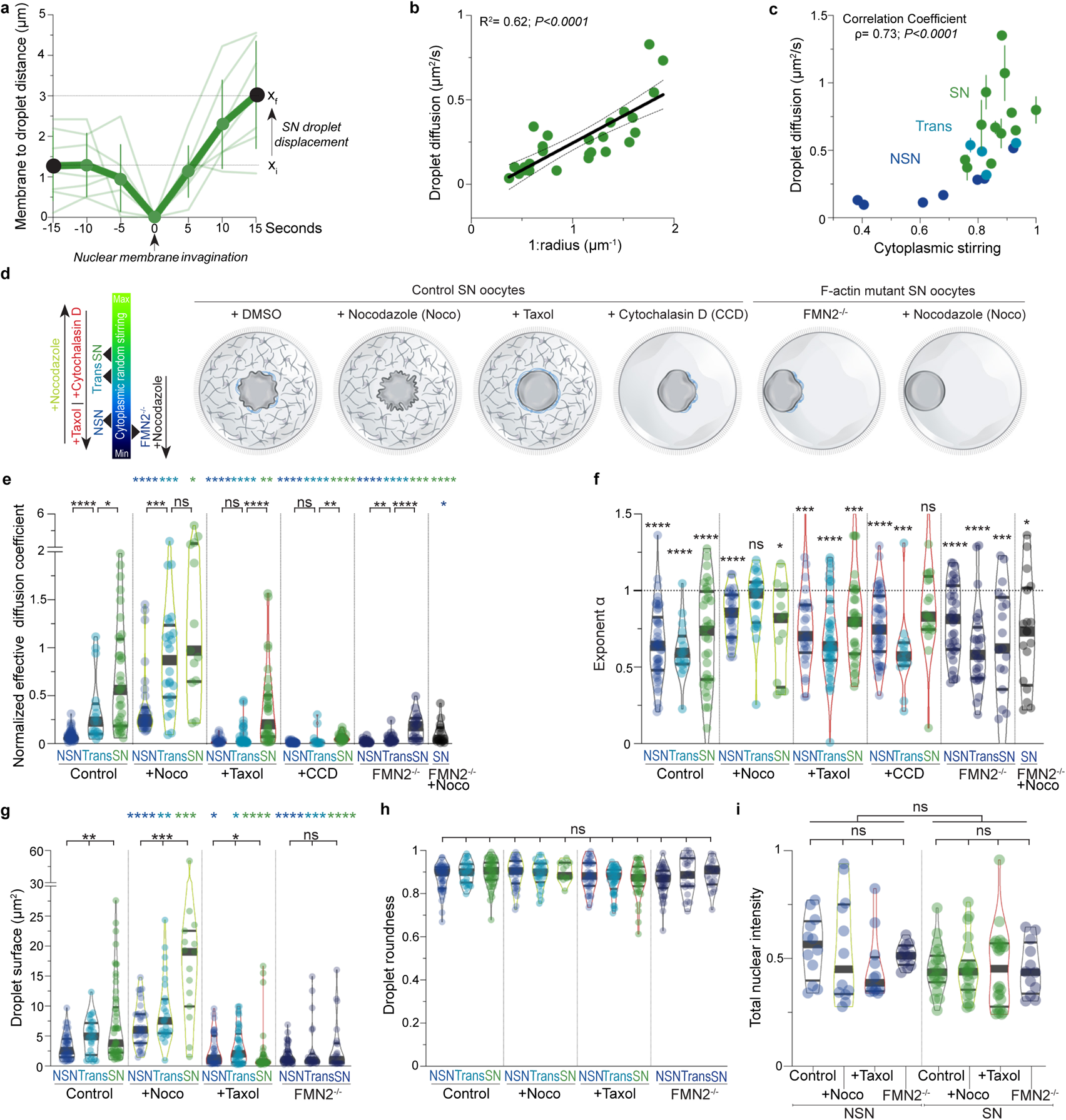
Cytoplasmic forces boost diffusive dynamics of nuclear SRSF2-GFP droplets on second to minute timescales. (**a**) Cytoplasmic forces locally displace SRSF2-GFP droplets via nuclear membrane fluctuations; graph shows measurements of the decreasing distance between the nuclear membrane’s invaginating tip and the SRSF2-GFP droplet that is pushed and displaced from x_i_ to x_f_; mean±s.d. of 7 droplets in 6 SN oocytes from films similar to the ones in Fig.2a and Supplementary Video 3. (**b**) Global SRSF2-GFP droplet diffusion inversely depends on droplet radius; 26 SN droplet diffusion coefficients plotted against the droplets’ radii^−1^ with a simple linear regression±90%CI; the relation is consistent with the *r^−1^* dependence expected from Stokes’ law. (**c**) Global diffusion of nuclear SRSF2-GFP droplets correlates with the intensification of cytoplasmic stirring that occurs during oocyte growth; mean±s.e.m. of 23 droplet diffusion coefficients plotted against cytoplasmic stirring values obtained using image correlation analyses; Spearman correlation coefficient H is shown. (**d**) Illustration of tools used to modulate cytoplasmic forces and consequent nucleus agitation in growing oocytes^7^; Nocodazole amplifies nucleus agitation whereas Taxol obstructs it; Cytochalasin-D disrupts the actomyosin network and thus nucleus agitation; mutant FMN2^−/−^ oocytes lack F-actin and only have residual microtubule-based agitation of the nucleus that is further diminished with the addition of Nocodazole; SN oocytes with actomyosin networks and microtubule organizing centers surrounding the nucleus are depicted when present, cytoplasmic microtubules are not illustrated. (**e-f**) Normalized effective diffusion coefficients of SRSF2-GFP droplets (**e**) and diffusive exponents α (**f**) in growing Control oocytes, oocytes incubated with Nocodazole, Taxol, or Cytochalasin-D, FMN2^−/−^ oocytes, and FMN2^−/−^ oocytes incubated with Nocodazole; effective diffusion coefficients were normalized by droplet size (*D_eff_* in 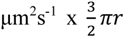 in µm); droplet number, Control NSN=46, Trans=17, SN=35, Nocodazole NSN=32, Trans=24, SN=13, Taxol NSN=29, Trans=43, SN=36, Cytochalasin-D NSN=48, Trans=14, SN=21, FMN2^−/−^ NSN=36, Trans=27, SN=19, FMN2^−/−^+Nocodazole SN=17. (**g-h**) Surface (**g**) and roundness (**h**) measurements of SRSF2-GFP droplets 3 hours post microinjection. Nocodazole and Taxol were added 2 hours post microinjection for an hour; droplet number, Control NSN=49, Trans=28, SN=58, Nocodazole NSN=38, Trans=28, SN=15, Taxol NSN=40, Trans=44, SN=39, FMN2^−/−^ NSN=50, Trans=26, SN=24. (**i**) Total nucleoplasmic SRSF2-GFP intensity measured on 40µm z-projections in NSN and SN Control oocytes, oocytes incubated with Nocodazole or Taxol, and FMN2^−/−^ oocytes; droplet number, Control NSN=12, SN=22, Nocodazole NSN=12, SN=22, Taxol NSN=12, SN=24, FMN2^−/−^ NSN=12, SN=12. Violin plots with median±quartiles; colored asterisks are statistical comparisons with Controls; *P* values derived from two-tailed Mann-Whitney *U*-Tests (**e-g**), Wilcoxon Signed-Rank Test (**f**; compared with a hypothetical I=1 median), and Kruskal-Wallis Tests (**g-i**), ns, not significant, *P*>0.0701, \**P*<0.0478, \*\**P*<0.0054, \*\*\**P*<0.001, \*\*\*\**P*<0.0001; oocyte growth color codes based on cytoplasmic stirring intensities.

**Extended Data Fig. 6.**
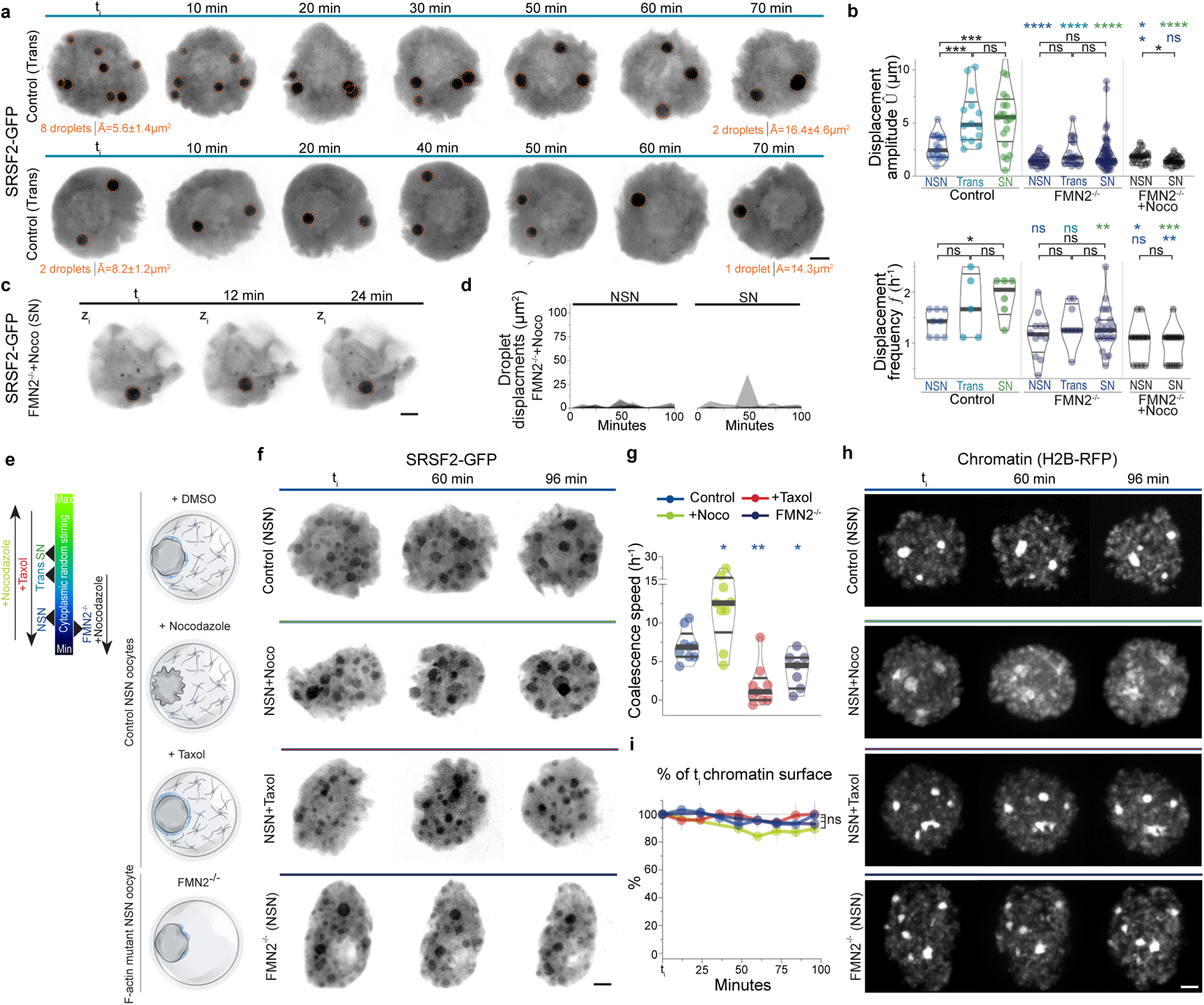
Cytoplasmic forces drive stochastic displacements and collision-coalescence of nuclear SRSF2-GFP droplets on minute to hour timescales. (**a**) Additional examples of Control SRSF2-GFP droplet minute-scale 3D-dynamics (50µm z-projections) in Trans cells highlighting random nucleoplasmic displacements and collision-coalescence dynamics; droplet number and mean surface evolution indicated below in orange. (**b**) Quantifications of displacement amplitudes (top) and frequencies (bottom) obtained with 3D-tracking of random SRSF2-GFP droplet displacements in nuclei of growing Control, FMN2^−/−^, and FMN2^−/−^+Nocodazole oocytes; number of amplitude measurements, Control NSN=19, Trans=14, SN=20, FMN2^−/−^ NSN=25, Trans=21, SN=57, FMN2^−/−^+Nocodazole NSN=20, SN=22; droplet number for frequency, Control NSN=8, Trans=5, SN=6, FMN2^−/−^ NSN=12, Trans=9, SN=23, FMN2^−/−^+Nocodazole NSN=11, SN=14; related to Fig.2g-h. (**c**) Time-lapse (single 0.5µm z-plane) showing weak minute-timescale displacements of a tracked SRSF2-GFP droplet in the nucleoplasm of a fully-grown SN FMN2^−/−^ oocyte incubated with Nocodazole. (**d**) 3D-tracks of SRSF2-GFP droplets in NSN and SN FMN2^−/−^ oocytes incubated with Nocodazole; droplet displacement amplitude Û and frequency *f* in b; droplet number, NSN=11, SN=14. (**e**) Illustration of tools used to modulate cytoplasmic forces in NSN oocytes for SRSF2-GFP droplet collision-coalescence experiments in f-h, Fig.2j and Supplementary Video 6; NSN oocytes with actomyosin networks and microtubule organizing centers surrounding the nucleus are depicted when present, cytoplasmic microtubules are not illustrated.(**f**) 3D-dynamics (40µm z-projections) of SRSF2-GFP droplets in NSN Control oocytes, oocytes incubated with Nocodazole or Taxol, and FMN2^−/−^ oocytes; droplet number and size evolution quantified in Fig.2j; see Supplementary Video 6. (**g**) Quantification of droplet coalescence speed of cells quantified in Fig.2j; Control n=9, Noco n=9, Taxol n=9, FMN2^−/−^ n=7. (**h**) Time-lapse images of chromatin (H2B-RFP; 40µm z-projections) showing absence of chromatin condensation in NSN oocytes analyzed in f and Fig.2j. (**i**) Quantification of chromatin surface evolution in % of initial surface; oocyte number, 4 per condition. Violin plots with median±quartiles (**b, g**); error bars represent mean±s.e.m. (**i**); colored asterisks in b are statistical comparisons with Control droplets; *P* values derived from two-tailed Mann-Whitney *U*-Tests (**b, g**) and Wilcoxon Matched-Pairs Signed Rank Tests (**i**), ns, not significant, *P*>0.0625, \**P*<0.042, \*\**P*<0.0079, \*\*\**P*<0.0008, \*\*\*\**P*<0.0001; oocyte growth color codes based on cytoplasmic stirring intensities and inhibitor treatment; scale bars, 5µm.

**Extended Data Fig. 7.**
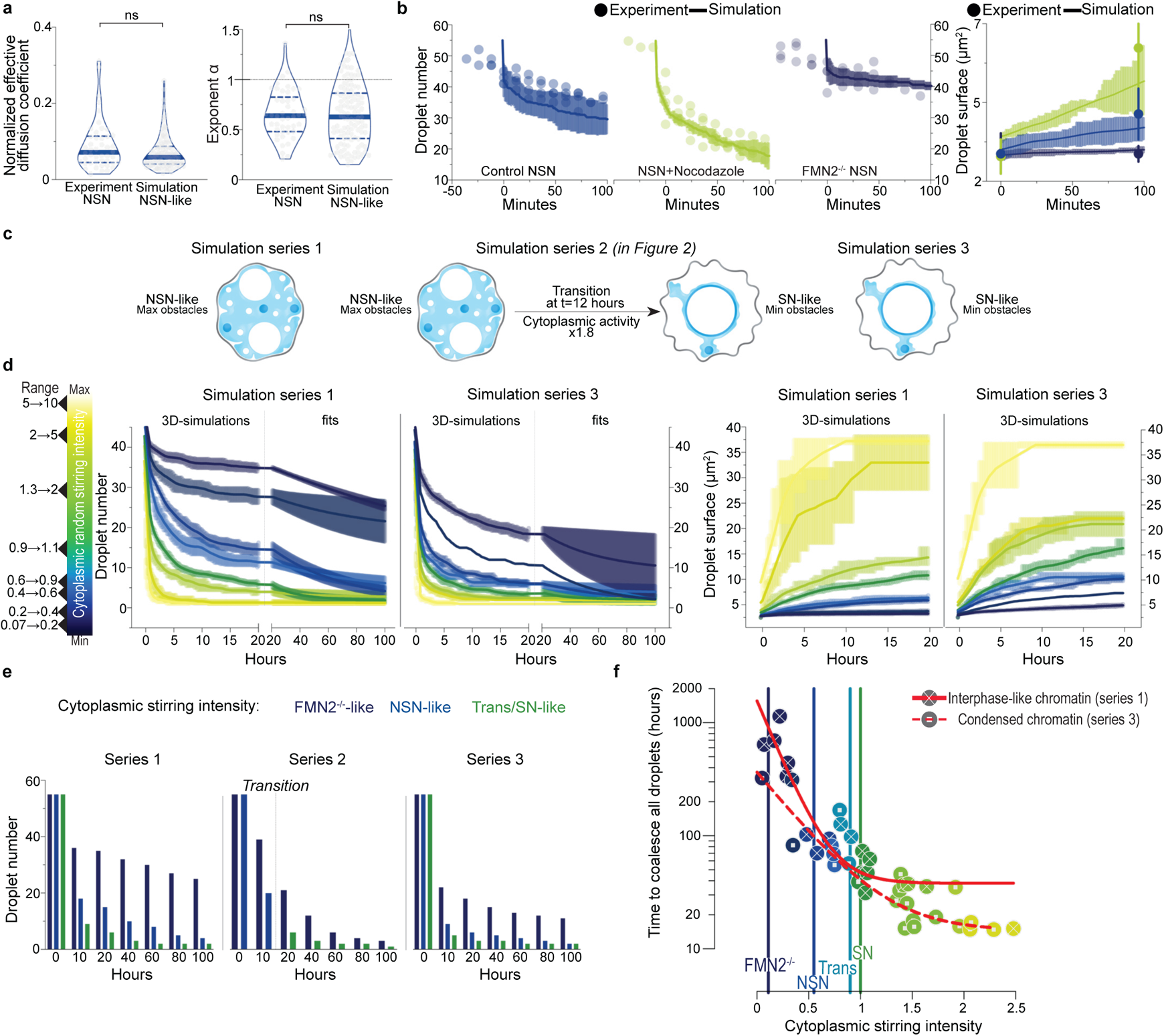
3D-simulations of nuclear SRSF2 droplet collision-coalescence dynamics relative to cytoplasmic forces on short (seconds) to long (days) timescales. (**a**) Effective diffusion coefficients normalized by droplet size (left) and droplet diffusive exponents I (right) from SRSF2-GFP experiments (NSN oocytes) and simulations with NSN-like obstacles; droplet number, Experiment=46; Simulation=115; violin plots with median±quartiles; *P* values derived from two-tailed Mann-Whitney *U*-Tests, ns, not significant, *P*=0.1203 and *P*=0.8741. (**b**) Droplet number (left) and size (right) evolution in NSN-like simulations (lines) and in NSN-oocyte experiments (circles) for Control, Nocodazole and FMN2^−/−^ conditions; simulations and experiments were aligned on comparable time frames by defining 45 droplets as t=0; simulations performed 5 times per condition and experimental data from 4 NSN oocytes per condition. (**c**) Schematic representation of the simulation regimes tested; Series 1: NSN-like configuration with interphase chromatin-like obstacles widely spread in the nucleus and cytoplasmic stirring activity maintained constant (e.g. 0.55 for Control, 1.05 for Nocodazole, and 0.11 for FMN2^−/−^); Series 2: NSN-to-SN-like configuration; first 12 hours of simulations are performed with the same parameters as in NSN-like simulations of series 1; nuclear obstacle and cytoplasmic stirring activity switch occurs at 12 hours whereby 40 % of chromatin-like obstacles surround the nucleolus and cytoplasmic activity is nearly doubled to mimic the transition into the SN-like condition, physiologically marked by chromatin condensation and cytoplasmic force intensification; Series 3: SN-like configuration with 40 % of chromatin-like obstacles surrounding the nucleolus and cytoplasmic stirring activity maintained constant (e.g. 1 for Control, 1.9 for Nocodazole, and 0.2 for FMN2^−/−^). (**d**) Droplet number (left) and size (right) evolution in NSN-like (series 1) and SN-like (series 3) simulations relative to varying cytoplasmic stirring activities; simulations are binned by range of activities (color gradient), from lower values (dark blue) to higher values (light yellow); droplet diffusion and fusions are simulated for 20 hours with longer-term dynamics predicted by a decreasing exponential fit calculated on the last 5h of the simulations; simulation number, 1^st^ series=44; 3^rd^ series=41. (**e**) Comparative histograms of droplet number evolution from the three simulations with cytoplasmic stirring activity intensities selected to resemble that of NSN and SN Control and F-actin mutant oocytes; note that in series 2, blue NSN-like bars switch to green Trans/SN-like bars after the transition. (**f**) Time necessary to coalesce 45 droplets relative to cytoplasmic stirring intensity comparing series 1 (NSN-like nucleus state with interphase-like chromatin) and 3 (SN-like nucleus state with condensed chromatin); time in the y-axis is calculated from fits in d; one-phase decay fits are in red and intersect vertical colored bars showing the simulated stirring activities corresponding to experimental ones in Controls and F-actin mutants; comparing plain and dashed intersection points allows to project the contributions of cytoplasmic activity versus chromatin to the droplet collision-coalescence drive in distinct conditions (e.g. Control/FMN2^−/−^) or stages within a condition (e.g. Control NSN/Trans/SN).

**Extended Data Fig. 8.**
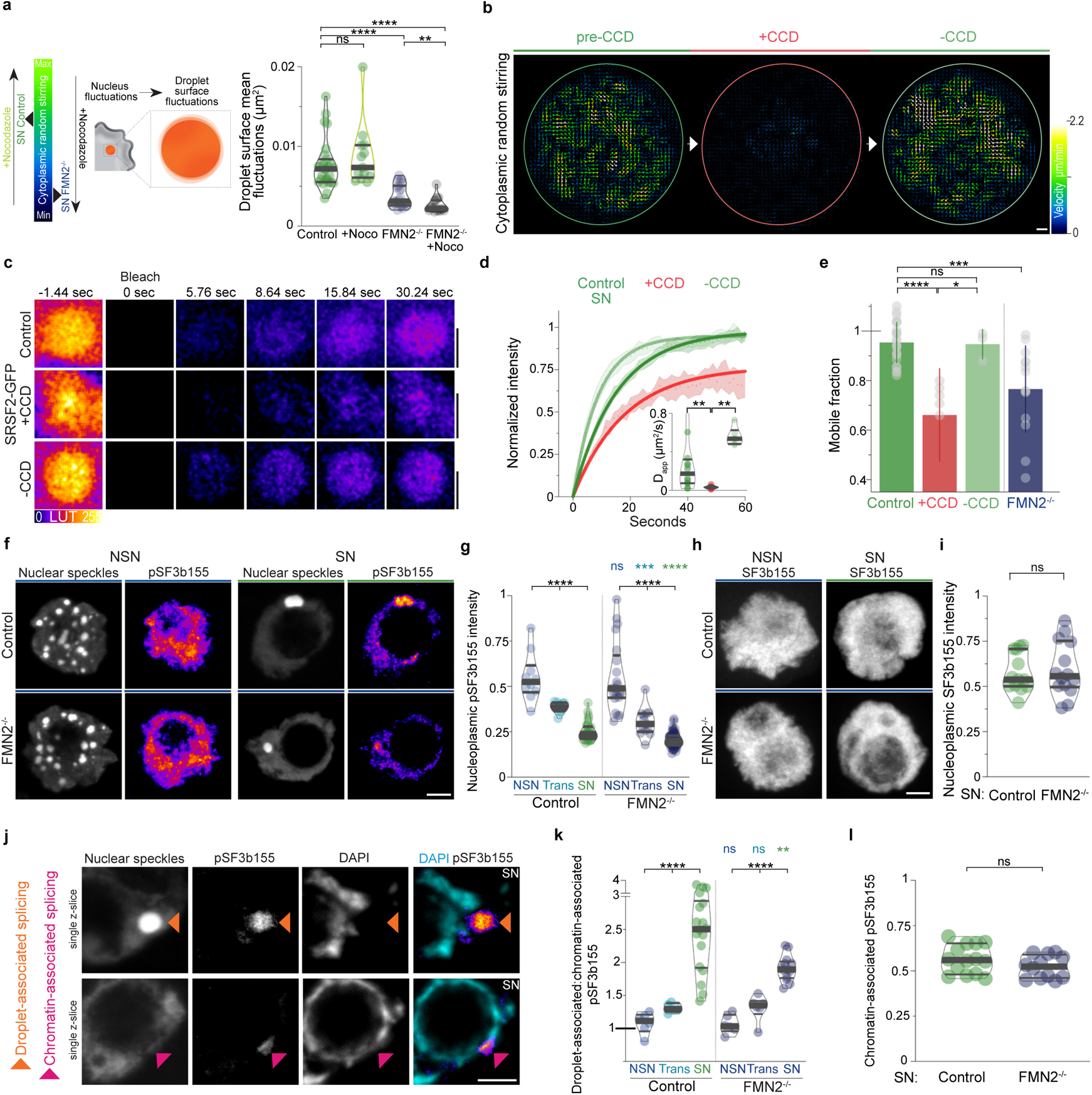
Cytoplasmic forces enhance nuclear speckle sub-droplet scale kinetics and mRNA splicing activity. (**a**) Cytoplasmic stirring drives droplet surface fluctuations; scheme of droplet surface fluctuations as a function of cytoplasmic stirring intensity (left); mean droplet surface fluctuations over 15 seconds (Δt=0.5s) in SN Control±Nocodazole and FMN2^−/−^±Nocodazole oocytes related to heatmaps in Fig.3a (right); comparably sized larger droplets were selected for analyses with a range of droplet radii from 2 to 2.7 µm; droplet number, Control=27; Nocodazole=13; FMN2^−/−^=23; FMN2^−/−^+Nocodazole=17. (**b**) Cytoplasmic stirring is restored after Cytochalasin-D washout; cytoplasmic flow vector maps generated by STICS analyses of bright-field 240 seconds-stream videos of a SN Control oocyte before Cytochalasin-D (left) that was incubated with Cytochalasin-D for 60 minutes (center; +CCD) prior to Cytochalasin-D washout for 60 minutes (right; −CCD); oocyte cortex is outlined; maps are color coded according to velocity magnitude. (**c**) Representative FRAP sequences of SRSF2-GFP droplets in SN Control oocytes, +CCD oocytes, and −CCD oocytes. (**d**) Normalized fluorescence intensity recovery curves (mean±s.e.m.) with simple exponential fits of SRSF2-GFP droplets in Control, Cytochalasin-D treated (+CCD), or Cytochalasin-D washout (-CCD) SN oocytes; insets, apparent diffusion coefficients (*D_app_*); droplet number, Control=12, +CCD=7, −CCD=6. (**e**) Mobile fractions of total populations for SRSF2-GFP droplets in SN Control oocytes, +CCD oocytes, −CCD oocytes, and FMN2^−/−^ oocytes; mobile fractions correspond to recovery fractions evaluated at curve plateaus fixed at 60 seconds for all conditions; droplet number, Control=24, +CCD=6, −CCD=3, FMN2^−/−^=14. (**f**) Representative co-immunostainings of nuclear speckles and the active spliceosome (engaged in mRNA splicing) marker pT313-SF3b155 in NSN (left) and SN (right) Control or FMN2^−/−^ oocytes (5µm z-projections); note the decrease of total nucleoplasmic active splicing with growth progression in both Control and mutant contexts that is consistent with the drop in transcription with growth progression; also note how active splicing becomes predominantly droplet-associated in both Controls and mutants during the NSN-to-SN transition. (**g**) Quantifications of global nucleoplasmic pT313-SF3b155 intensities in Control and FMN2^−/−^ oocytes at the three growth stages; cell number, Control NSN=10, Trans=11, SN=32, FMN2^−/−^ NSN=21, Trans=13, SN=64. (**h**) Representative immunostainings of the non-phosphorylated form of SF3b155 showing comparable nucleoplasmic intensity levels in NSN and SN Control or FMN2^−/−^ oocytes (35µm z-projections). (**i**) Quantifications of total nucleoplasmic SF3b155 intensity normalized by cytoplasmic noise in SN Control and FMN2^−/−^ oocytes; cell number, Control=13, FMN2^−/−^=14. (**j**) Representative single z-sections of co-immunostainings of active splicing (pT313-SF3b155) associated with nuclear speckles (orange arrowhead) or with DNA (fuchsia arrowhead) in SN oocytes. (**k**) Quantifications of droplet-associated to chromatin-associated active splicing (pT313-SF3b155) signal ratios in Control and FMN2^−/−^ oocytes at the three growth stages; cell number, Control NSN=6, Trans=5, SN=18, FMN2^−/−^ NSN=6, Trans=6, SN=13. (**l**) Quantifications of chromatin-associated active splicing (pT313-SF3b155) in 14 Control and 12 FMN2^−/−^ SN oocytes. Violin plots with median±quartiles (**a, d, g, i, k-l**); error represents mean±s.d. (**e**); *P* values derived from two-tailed Mann-Whitney *U*-Tests (**a, d, e, i, k-l**) and from Kruskal-Wallis Tests (**g**), ns, not significant, *P*>0.3307, \**P*=0.0238, \*\**P*<0.0083, \*\*\**P*<0.0005, \*\*\*\**P*<0.0001; scale bars, (**c**) 3µm and (**b, f, h, j**) 5µm.

**Extended Data Fig. 9.**
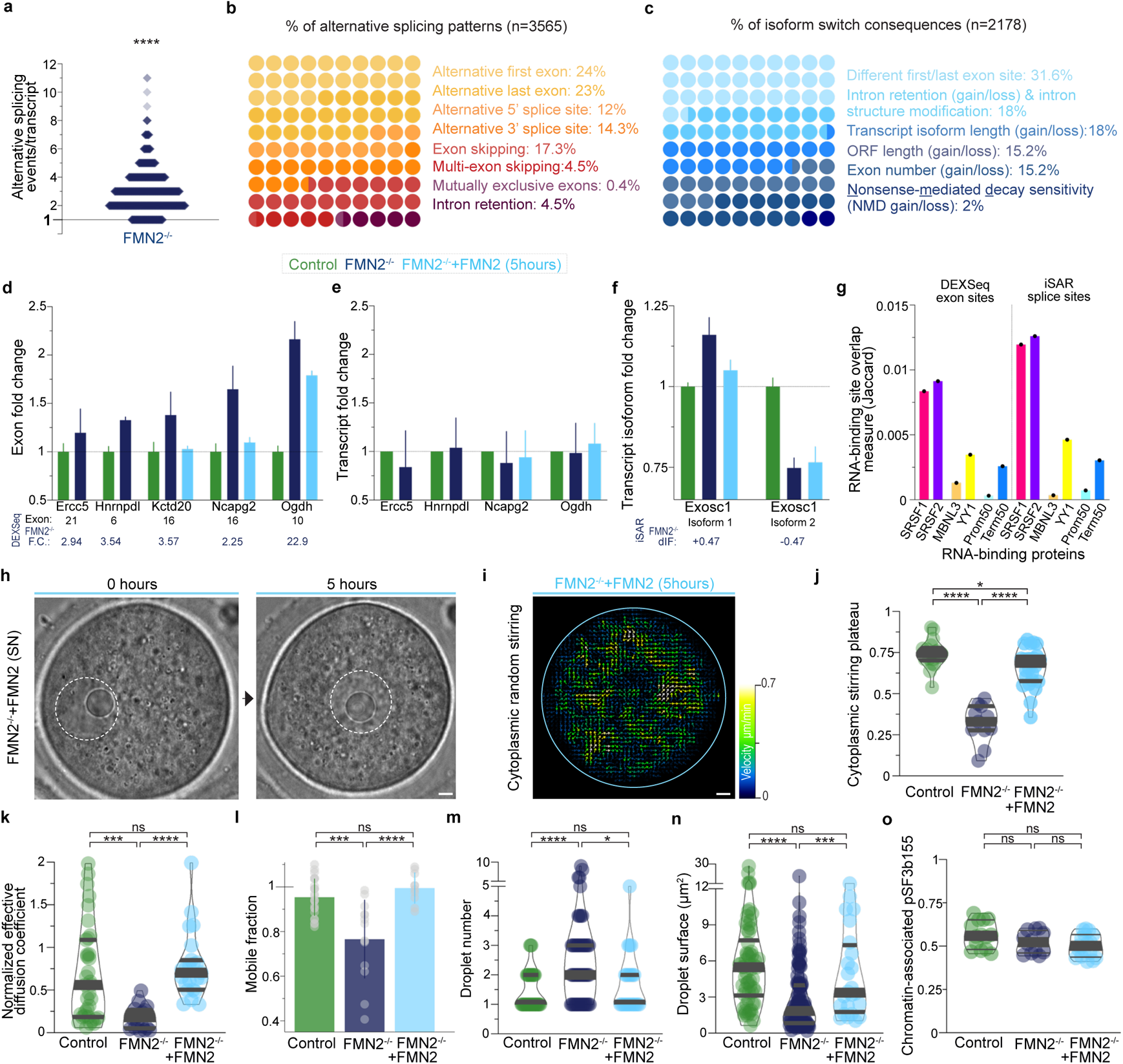
Cytoplasmic force disruption alters mRNA splicing; restoring cytoplasmic forces in mutant SN oocytes fully rescues droplet number, size, and multiscale dynamics along with a partial reversal of mRNA splicing pattern alterations within 5 hours. (**a**) Multiple alternative splicing events occur per transcript in FMN2^−/−^ oocytes with a median of 3; each plotted dot corresponds to one or more alternative splicing events detected in a single individual transcript for a total of 1259 transcripts. (**b**) 10×10 dot plot showing the percentages of mRNA alternative splicing events detected in FMN2^−/−^ (n=3565 events) and color-coded according to categories of alternative splicing event types; see Supplementary Table 2. (**c**) 10×10 dot plot showing the percentages of predicted consequences of mRNA isoform switches in FMN2^−/−^ (n=2178 consequences) and color-coded according to consequence categories; see Supplementary Table 2. (**d-e**) RT-qPCR validation of FMN2^−/−^ differential exon usage in distinct genes obtained bioinformatically with DEXSeq (Supplementary Table 1) and partial reversal of tendencies after expression of FMN2 in FMN2^−/−^ oocytes for 5 hours; exon quantities are in (**d**) and associated transcript quantities are in (**e**); results are averaged from three independent experiments (mean±s.e.m.) with heterozygous FMN2^+/−^ as Controls; FMN2^−/−^ exon usage tendencies were confirmed (compare with DEXSeq fold change values below); note that exon numbers shown were defined by DEXSeq. (**f**) RT-qPCR validation of FMN2^−/−^ alternative splicing detected bioinformatically with IsoformSwitchAnalyzeR (Supplementary Table 2) and partial reversal of tendencies after expression of FMN2 in FMN2^−/−^ oocytes for 5 hours; results are averaged from three independent experiments (mean±s.e.m.) with heterozygous FMN2^+/−^ as Controls; FMN2^−/−^ alternative splicing tendencies were confirmed in Exosc1 transcript isoforms (compare with dIF values below). (**g**) Jaccard measures of overlaps between sets of genomic sites, through computation of the ratio of their intersections to their union^75^; differentially used exon sites (DEXSeq) and alternative splice sites of transcripts (iSAR) of FMN2^−/−^ oocytes were tested bidirectionally for spatial correlation with SRSF1, SRSF2, MBNL3, and YY1 RNA-binding sites^28–30^, as well as against *in silico* controls corresponding to the 50 first (Prom50) or last (Term50) nucleotides of all RefSeqNCBI transcripts; all Jaccard measures were significant (*P*<0.02) except for the overlap between differentially used exon sites and Prom50 sites; see Supplementary Table 3. (**h**) Time-lapse of an FMN2^−/−^ SN oocyte microinjected with FMN2 cRNA showing the repositioning of the nucleus within 5 hours. (**i**) Cytoplasmic stirring in FMN2^−/−^ SN oocytes is restored after FMN2 expression; cytoplasmic flow vector map generated by STICS analyses of a bright-field 120 seconds-stream video of a SN FMN2^−/−^ oocyte 5 hours after microinjection with FMN2 cRNA; oocyte cortex is outlined; maps are color coded according to velocity magnitude. (**j**) Cytoplasmic stirring intensity plateaus in SN Control, FMN2^−/−^, and FMN2^−/−^+FMN2 (5hours) oocytes obtained with image correlation analyses of cytoplasmic stirring in live oocytes for 100 seconds; cell number, Control=18, FMN2^−/−^=11, FMN2^−/−^+FMN2=24. (**k**) Normalized effective diffusion coefficients of SRSF2-GFP droplets in SN Control, FMN2^−/−^, and FMN2^−/−^+FMN2 (5hours) oocytes; effective diffusion coefficients were normalized by droplet size (*D_eff_* in 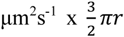 in µm); droplet number, Control=35, FMN2^−/−^ =18, FMN2^−/−^+FMN2=23. (**l**) Mobile fractions of total populations for SRSF2-GFP droplets in SN Control, FMN2^−/−^, and FMN2^−/−^+FMN2 (5hours) oocytes; mobile fractions correspond to recovery fractions evaluated at curve plateaus fixed at 60 seconds for all conditions; droplet number, Control=24, FMN2^−/−^=14, FMN2^−/−^+FMN2=10; related to Fig.3b-c. (**m-n**) Quantifications of nuclear speckle droplet number (**m**) and surface (**n**) in nuclei of fully grown SN Control, FMN2^−/−^, and FMN2^−/−^+FMN2 (5hours) oocytes; speckles counted in 41 Control, 52 FMN2^−/−^, and 24 FMN2^−/−^+FMN2 cells, measured 62 Control, 112 FMN2^−/−^, and 24 FMN2^−/−^+FMN2 droplets. (**o**) Quantifications of chromatin-associated active splicing (pT313-SF3b155) in 14 Control, 12 FMN2^−/−^ and 17 FMN2^−/−^+FMN2 SN oocytes. Violin plots with median±quartiles (**j-o**); *P* values derived from Wilcoxon Signed-Rank Test against a hypothetical median of 1(**a**) and two-tailed Mann-Whitney *U*-Tests (**j-o**), ns, not significant, *P*>0.055, \**P*<0.0233, \*\*\**P*<0.0009, \*\*\*\**P*<0.0001; scale bars, 5µm.

**Extended Data Fig. 10.**
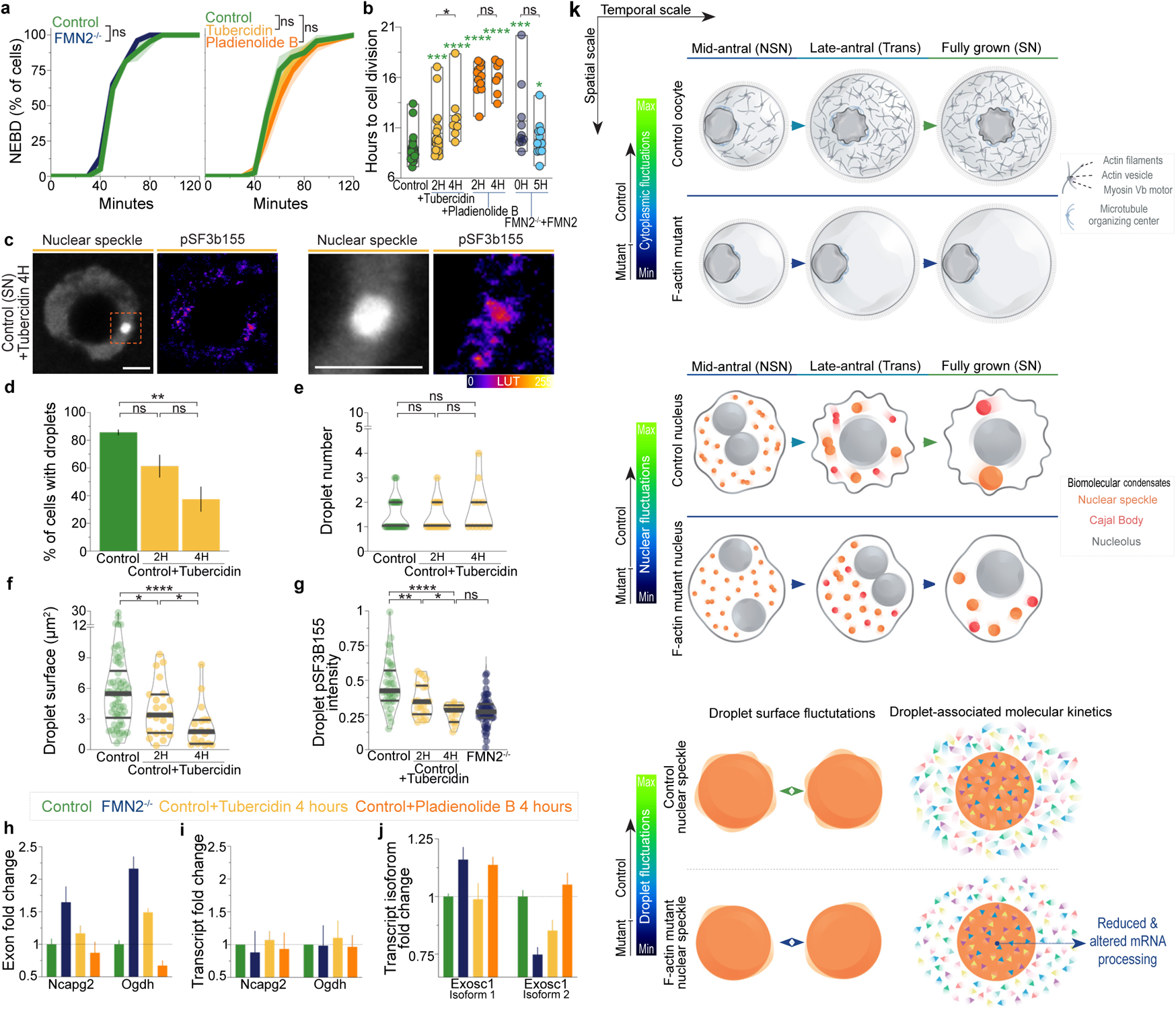
Consequences of splicing modulation in SN oocytes; model with cytoplasmic stirring forces as scale-crossing organizers of nuclear condensates in growing oocytes. (**a**) Timing of nuclear envelope breakdown (NEBD) in percent of cells for Control and FMN2^−/−^ SN oocytes (left) or for Control, Control+Tubercidin (2-4 hours), and Control+Pladieonolide B (2-4 hours) oocytes (right); SN oocytes were washed 6 times in fresh M2+BSA medium to remove traces of Milrinone and splicing modulators before tracking NEBD every 10 minutes for 2 hours; cell number, Control=20, FMN2^−/−^=53, Control=60, Control+Tubercidin 2-4H=78, Control+Pladieonolide B 2H-4H=70. (**b**) Time necessary in hours to complete the meiotic division for oocytes that did divide after treatments with splicing modulators (2-4 hours of Tubercidin or Pladienolide B) or FMN2-rescues (0-5 hour incubation after microinjection) at the SN stage; cells were filmed every 3 minutes up to 21 hours; NEBD defined the start point and polar body extrusion defined the timing of cell division; cell number, Control=27, Control+Tubercidin 2H=18, 4H=8, Control+Pladieonolide B 2H=13, 4H=7, FMN2^−/−^+FMN2 0H=10, 5H=12. (**c**) Representative co-immunostainings of nuclear speckles and the active spliceosome (engaged in mRNA splicing) marker pT313-SF3b155 in SN Control oocytes treated for 4 hours with Tubercidin (5µm z-projections); dashed orange box indicates the magnification shown on the right. (**d**) Proportions of SN Control oocytes±Tubercidin (2 to 4 hours) that contain nuclear speckle droplets in their nucleus; cell number, Control=54, Control+Tubericidin 2H=27,4H=16. (**e**) Quantifications of nuclear speckle droplet number in SN Control oocytes±Tubercidin (2 to 4 hours); cell number, Control=41, Control+Tubericidin 2H=16,4H=11. (**f**) Quantifications of nuclear speckle droplet surface in SN Control oocytes±Tubercidin (2 to 4 hours); droplet number, Control=62, Control+Tubericidin 2H=22,4H=18. (**g**) Quantifications of single nuclear speckle droplet-associated active splicing (pT313-SF3b155) intensities in SN Control oocytes±Tubercidin (2 to 4 hours); droplet number, Control=38, Control+Tubericidin 2H=21,4H=15. (**h-j**) Assessment of alternative splicing tendencies in SN oocytes using RT-qPCR of Control+Tubercidin (4 hours) and Control+Pladieonolide B (4 hours) cells relative to Control and FMN2^−/−^ cells; exon quantities are in (**h**) and associated transcript quantities are in (**i**); alternative splicing tendencies were assessed for Exosc1 transcript isoforms (**j**); results are averaged from three independent experiments (mean±s.e.m.); related to Extended Data Fig.9d-f. Floating bars with mean±range (**b**), histogram with mean±s.e.m. (**d**), violin plots with median±quartiles (**e-g)**; *P* values derived from *P* values derived from two-tailed Wilcoxon matched-pairs signed rank tests (**a**) and two-tailed Mann-Whitney *U*-Tests (**b, d-g**), ns, not significant, *P*>0.0577, \**P*<0.0278,\*\**P*<0.0088,\*\*\**P*<0.0005, \*\*\*\**P*<0.0001; scale bars, 5µm. (**k**) Mouse oocyte growth model with cytoplasmic random stirring forces as multiscale and functional reorganizers of nuclear droplets. At the cell scale (top), the intensification of cytoplasmic stirring nonspecifically transports the nucleus over tens of microns while rapidly agitating it^5,7,18^. At the nucleus scale (center), the intensification of cytoplasmic stirring orchestrates reorganization of nuclear compartments, by accelerating diffusive dynamics and collision-coalescence of liquid biomolecular condensates. At the droplet scale (bottom), cytoplasmic stirring drives nuclear droplet surface fluctuations and boosts droplet-associated molecular kinetics, enhancing condensate-associated biomolecular reactions and promoting meiotic success. Inversely, cells with diminished cytoplasmic stirring due to cytoskeletal defects present organizational flaws associated with cell-, nucleus-, and droplet-scale dynamics. Weak cytoplasmic stirring like in F-actin mutants is insufficient to displace the nucleus or drive proper multiscale reorganization of nuclear liquid condensates. This multiscale disorganization leads to droplet-associated alterations in the processing of mRNA and major defects in subsequent oocyte development that drives fertility. Thus, cells can deploy cytoplasmic forces to functionally refashion liquid compartments in membrane-bound organelles like the nucleus for developmental success.

## Supplementary Video 1-7 Legends

**Video 1 | Live-imaging of cytoplasmic random stirring in growing Control and fully-grown Control or F-actin mutant oocytes**

Time-lapse stream-mode imaging in bright-field of a Control growing (NSN from a mid-antral follicle; left) oocyte and fully-grown Control and FMN2^−/−^ mouse oocytes (SN; center and right) showing the increase in short-timescale cytoplasmic stirring with growth and disrupted activity in the fully-grown F-actin mutant; note the peripheral nuclei in Control NSN and FMN2^−/−^cells and the presence of 2 visible nucleoli in the NSN nucleus. See Fig.1 and Extended Data Fig.1. Time in mm:ss; scale bar, 5µm.

**Video 2 | Live-imaging of SRSF2-GFP droplet coalescence in the fully-grown oocyte nucleus**

Time-lapse stream-mode imaging of two nucleoplasmic SRSF2-GFP droplets fusing in an SN oocyte; SRSF2 is a nuclear speckle marker. See Extended Data Fig.3. Time in mm:ss; scale bar, 5µm.

**Video 3 | Live-imaging of local nuclear SRSF2-GFP droplet displacements and collision-coalescence driven by cytoplasmic stirring in growing oocytes**

Time-lapse stream-mode imaging of nucleoplasmic SRSF2-GFP droplets physically pushed by cytoplasm-based random kicks of the nuclear membrane in NSN and SN oocytes; droplet collision-coalescence is instigated by the cytoplasm-based forces that locally displace the NSN droplet; note the more rapid diffusive dynamics of the SN droplet. See Fig.2 and Extended Data Fig.5. Time in mm:ss; scale bar, 5µm.

**Video 4 | Live-imaging of global nuclear SRSF2-GFP droplet dynamics in growing oocytes with Control or modulated cytoplasmic forces**

Time-lapse stream-mode imaging of global SRSF2-GFP droplet diffusive dynamics in nucleoplasms of NSN (1^st^ row) and SN (2^nd^ row) oocytes with Control cytoplasmic forces (1^st^ column), amplified forces (+Nocodazole; 2^nd^ column), obstructed forces (+Taxol; 3^rd^ column), and disrupted forces (FMN2^−/−^; 4^th^ column). See Fig.2 and Extended Data Fig.5. Time in mm:ss; scale bar, 5µm.

**Video 5 | Live-imaging of nuclear SRSF2-GFP droplets with random large-scale displacements and collision-coalescence in growing oocytes**

Time-lapse images (50 µm z-projections) of global nucleoplasmic SRSF2-GFP droplet dynamics in Trans and SN oocytes on longer timescales; note random large-scale displacements of droplets and occasional collision-coalescence. See Fig.2 and Extended Data Fig.6. Time in hh:mm; scale bar, 5µm.

**Video 6 | Live-imaging of nuclear SRSF2-GFP droplet coalescence dynamics in growing oocytes with Control or modulated cytoplasmic forces**

Time-lapse images (40 µm z-projections) of nucleoplasmic SRSF2-GFP droplet dynamics showing collision-coalescence evolution on longer timescales in NSN oocytes with Control cytoplasmic forces (1^st^ column), amplified forces (+Nocodazole; 2^nd^ column), obstructed forces (+Taxol; 3^rd^ column), and disrupted forces (FMN2^−/−^; 4^th^ column); note that starting point droplet numbers for each condition vary slightly. See Fig.2 and Extended Data Fig.6. Time in hh:mm; scale bar, 5µm.

**Video 7 | 3D-simulations of nuclear SRSF2 droplet collision-coalescence evolution from NSN-like to SN-like states relative to cytoplasmic force intensity**

Time-lapse images of 3D-simulations on hour to day timescales showing the evolution of nuclear SRSF2-like droplet collision-coalescence relative to varying intensities of cytoplasmic stirring activity; NSN-to-SN-like simulation regime with first 12 hours of simulations performed with NSN-like parameters before a nuclear obstacle and cytoplasmic activity switch that occurs at 12 hours whereby 40 % of chromatin-like obstacles surround the nucleolus and cytoplasmic activity nearly doubles to mimic the transition into the SN-like condition (physiologically marked by chromatin condensation and cytoplasmic force intensification). FMN2^−/−^-like cytoplasmic forces (0.11-0.19; 1^st^ column), Control-like forces (0.55-1; 2^nd^ column), ~two-fold amplified forces (1.05-1.89; Control+Nocodazole-like; 3^rd^ column), and ~four-fold amplified forces (2.32-4.18; 4^th^ column). SRSF2 droplets are in orange and the nucleolus is in light grey placed in a dark grey spherical nucleus; chromatin-like obstacles are invisible. See Fig.2 and Extended Data Fig.7. Time in hh:mm; scale bar, 5µm.

## Supplementary Table 1-5 Legends

**Supplementary Table 1 | mRNA exon usage table (FMN2+/− vs. FMN2−/− SN oocytes)**

Sheet 1, DEXSeq output with *P_adj_*<0.05. Sheet 2, Illustration summarizing primer pair design for RT-qPCR experiments. See Fig. 3 and Extended Data Fig.9.

**Supplementary Table 2 | mRNA isoform usage table (FMN2+/− vs. FMN2−/− SN oocytes)**

Sheet 1, IsoformSwitchAnalyzeR (iSAR) output with *P_adj_*<0.05. Sheet 2, detected alternative splicing events. Sheet 3, transcript ID and splicing coordinates. Sheet 4, Predicted isoform switch consequences. Sheet 5, isoforms selected for RT-qPCR validation and illustration of primer pair design. See Fig. 3 and Extended Data Fig.9.

**Supplementary Table 3 | Enrichment and spatial correlation analyses of altered splicing sites relative to SRSF1 and SRSF2 binding sites**

Sheet 1-2, Enrichment analyses of SRSF1 and SRSF2 binding sites^28^ in FMN2^−/−^ genes associated with altered splicing relative to all genomic transcripts; statistical tests are displayed in sheet 2. Sheet 3-4, Spatial correlation analyses using the GenometriCorr package^75^ comparing FMN2^−/−^ altered splicing sites with binding sites of SRSF1, SRSF2, MBNL3, or YY1(^28–30^); *in silico* controls correspond to the 50 first (Prom50) or last (Term50) nucleotides of all RefSeqNCBI transcripts; see Extended Data Fig.9.

**Supplementary Table 4 | Functional enrichment of FMN2−/− genes with mRNA processing alterations**

Enrichr Gene Ontology – Biological Process output for the list of FMN2^−/−^ genes affected by mRNA processing defects revealed by DEXSeq and iSAR.

**Supplementary Table 5 | FMN2−/− genes with mRNA processing alterations compared to translated ones during the first meiotic division**

FMN2^−/−^ genes with altered mRNA processing compared to translational status of transcripts during the first meiotic division^31^. Sheet 1, list of translated, activated (engaged in translation), and repressed (degraded) transcripts during the first meiotic division; GeneID MGI-conversions are next to each list. Sheet 2, Comparative analysis showing that 53% of FMN2^−/−^ genes with altered mRNA processing (DEXSeq and iSAR lists) are translated or engaged in translation during the first meiotic division.

## Notes

### Competing Interest Statement

The authors have declared no competing interest.

